# A large, distinct family of RNase H fold nucleases associated with numerous defense systems in prokaryotes

**DOI:** 10.64898/2026.11.03.763955

**Authors:** Yuri I. Wolf, Kanza Arshad, Kwame Kannatey-Asibu, Rachel Q. Theisen, Katherine A. Kitzinger, Preeti V. Jain, Eugene V. Koonin, Kira S. Makarova, Hannah E. Ledvina

**Affiliations:** Computational Biology Branch, Division of Intramural Research, National Library of Medicine, National Institutes of Health, Bethesda, MD 20894, USA; Department of Molecular, Cellular and Developmental Biology, University of Michigan, Ann Arbor, MI 48109, USA

## Abstract

During the last decade, systematic efforts aimed at the characterization of prokaryotic immunity resulted in the discovery of several hundred distinct defense systems in bacteria and archaea. This suggests that the common defense systems were already known and only rare ones remained to be identified. Here we present a genomic census and analysis of a vast family of bacterial and archaeal proteins (known as Domain of Unknown Function, DUF3800) that contain a distinct variant of the RNase H fold and are encoded in a striking 15% of the available complete prokaryotic genomes. Most of these proteins are encoded in defense islands, often in a close association with known defense systems, hence the designation WARD (<u>W</u>eaponized <u>A</u>ntiphage <u>R</u>Nase-dependent <u>D</u>efense). The WARDs display a broad variety of protein domain architectures and genomic contexts, including a strong link to restriction-modification, CRISPR-Cas, retrons, and non-coding RNAs, suggestive of a substantial diversity in defense mechanisms. We experimentally tested 10 WARDs, mostly, encoded by standalone genes, and found that 9 of these conferred robust protection against diverse phages in *Escherichia coli*. Therefore, WARDs appear to represent the third most abundant defense system type in prokaryotes.

## Introduction

Coevolution between viruses and their hosts generally follows the Red Queen evolutionary dynamic, yielding a multitude of diverse immune systems and mechanisms, while the viruses continuously evolve countermeasures to overcome these defenses ^1–4^. This arms race often results in the emergence of multilayer defenses whereby organisms activate different immune mechanisms synergistically ^5–10^. The first line of protection typically involves passive mechanisms of parasite avoidance, such as cell surface modification preventing virus entry ^8,11^. If the virus clears the first barrier and the organism is infected, innate or adaptive immunity pathways are activated that block virus replication via a broad variety of mechanisms, and if active immunity fails, the last resort defense is triggered, namely, programmed self-damage mechanisms, also known as abortive infection ^11–13^. In prokaryotes, active immunity pathway that clear viruses and self-damage pathways are often genetically linked and co-regulated. This phenomenon was initially predicted by comparative genomics ^6^, and subsequent research revealed an enormous variety and complexity of coordinated responses to infection ^14^.

Nucleases cleaving either DNA or RNA are the most common effectors in both innate and adaptive immunity, and abortive infection systems ^15,16^. Due to the constant pressure from viral antidefense factors, often targeting effectors, the latter tend to evolve fast and are frequently exchanged between defense systems of the same type. Numerous cases of effector shuffling have been described in restriction-modification (RM), CRISPR-Cas and other defense systems ^3,17–19^. In addition to their canonical roles as defense effectors, some of these nucleases can function as independent defense modules, as ancillary components of other systems, toxins in toxin-antitoxin (TA) modules, effectors of interspecies conflict systems, and also outside of the defense scope, being involved in DNA or RNA replication, recombination and repair ^15,16,20–22^. The PD-[DE]xK, RelE, PIN, HEPN, and Endo VII/HNH families are among the most abundant, versatile nucleases found in most of these contexts ^15,23–26^. RNase H family nucleases are well-studied, essential components of the prokaryotic DNA replication machinery that are responsible for the removal of RNA primers from the replicated DNA ^27,28^. RNase H fold nucleases are less common in defense but function as effectors in such well-characterized RNA-guided systems as Argonauts as well as type II and type V CRISPR-Cas (Cas9 and Cas12 proteins, respectively) ^29,30^.

The DUF3800 family, which also consists of (predicted) RNase H fold nucleases, has so far escaped close attention. The genes of this family have been spotted in the genomic contexts of established defense systems, such as CRISPR-Cas and retrons, but their functions have not been studied ^31–33^. An early study, published before DUF3800 proteins were predicted to possess nuclease activity, showed that one of these proteins, AbiI, encoded on a lactococcal plasmid provided defense against two lactococcal phages ^34^. The only other bit of experimental evidence is that one of the DUF3800 proteins is the toxin in a type I toxin-antitoxin (TA) system, in which the toxin mRNA translation is blocked by an antisense RNA antitoxin ^35^. Additionally, it has been found that DUF3800 proteins are often encoded next to a small, conserved non-coding (nc) RNA, suggesting that the two are functionally linked. Eleven distinct families of such DUF3800-associated RNAs have been predicted, but there is no experimental data on their functions ^36^. One possible role of the ncRNA co-located with DUF3800 genes might be the antitoxin function in type III TA modules. There are only a few known TA systems of this type, with toxins belonging to two distinct nuclease superfamilies, namely, ToxN/AbiQ/TepnN of the MazF RNase superfamily, and AbiF and AbiV of the HEPN RNase superfamily ^37–41^. In these systems, the ncRNA, typically folded into an elaborate structure, is encoded upstream of the toxin gene and serves as the antitoxin blocking the toxin activity. All known type III TAs are involved in antivirus defense ^38,40,41^.

Due to the high sequence divergence, DUF3800 domain-containing proteins are often overlooked or misannotated in public databases. Here we present a comprehensive census of the DUF3800 family and DUF8203 family sharing the same core fold with DUF3800 in bacterial and archaeal genomes, and the results of extensive computational analysis of these families, yielding insights into their organization, structure, possible functions and evolution. We provide initial experimental evidence that most of the DUF3800 proteins are components of diverse, novel defense systems.

## Results and Discussion

### DUF3800-containing proteins

To obtain a representative collection of DUF3800 and DUF8203 protein families, we used iterative sequence search against the NCBI clustered NR database using PSI-BLAST, followed by manual curation of sequences retrieved in each iteration (see Materials and Methods). The resulting validated set of the members of these two families was used to construct 500 sequence profiles covering the DUF3800 core domains and one for DUF8203 (Datasets 1 and 2). These profiles were used as queries for a second round of search of the clustered NR database using PSI-BLAST. The resulting set of retrieved sequences was combined with the previously identified ones and once again verified manually. Altogether, we identified 36,094 DUF3800 cluster representatives corresponding to 58,373 non-redundant proteins, and 152 DUF8203 cluster representatives, corresponding to 180 non-redundant proteins (Table S1 and S2).

DUF3800 homologs are present in both bacteria and archaea (95% and 5% of the identified family members, respectively) (Figure 1A). A small fraction (<1%) was also identified in bacterial viruses. Although two sequences were annotated as belonging to *Porifera* (eukaryotes), the lack of introns and the absence of genes of clear eukaryotic origin in the respective contigs suggests bacterial contamination (Table S1). To assess the prevalence of DUF3800 accounting for the sampling bias, we identified the genes of this family in a database of complete prokaryotic genomes (Dataset 3). Overall, DUF3800 family is represented in ∼15% of prokaryotic genomes, with paralogs detected in many cases (1.4 family members per genome, on average). DUF3800 proteins are conspicuously absent in *Chlamydiota* and are rare in *Campylobacterota*, *Fusobacteriota* and *Thermotogota*, whereas *Actinomycetota*, *Deinococcota* and *Chloroflexota* include the largest fractions of genomes encoding members of this family (Figure 1B).

**Figure 1.**
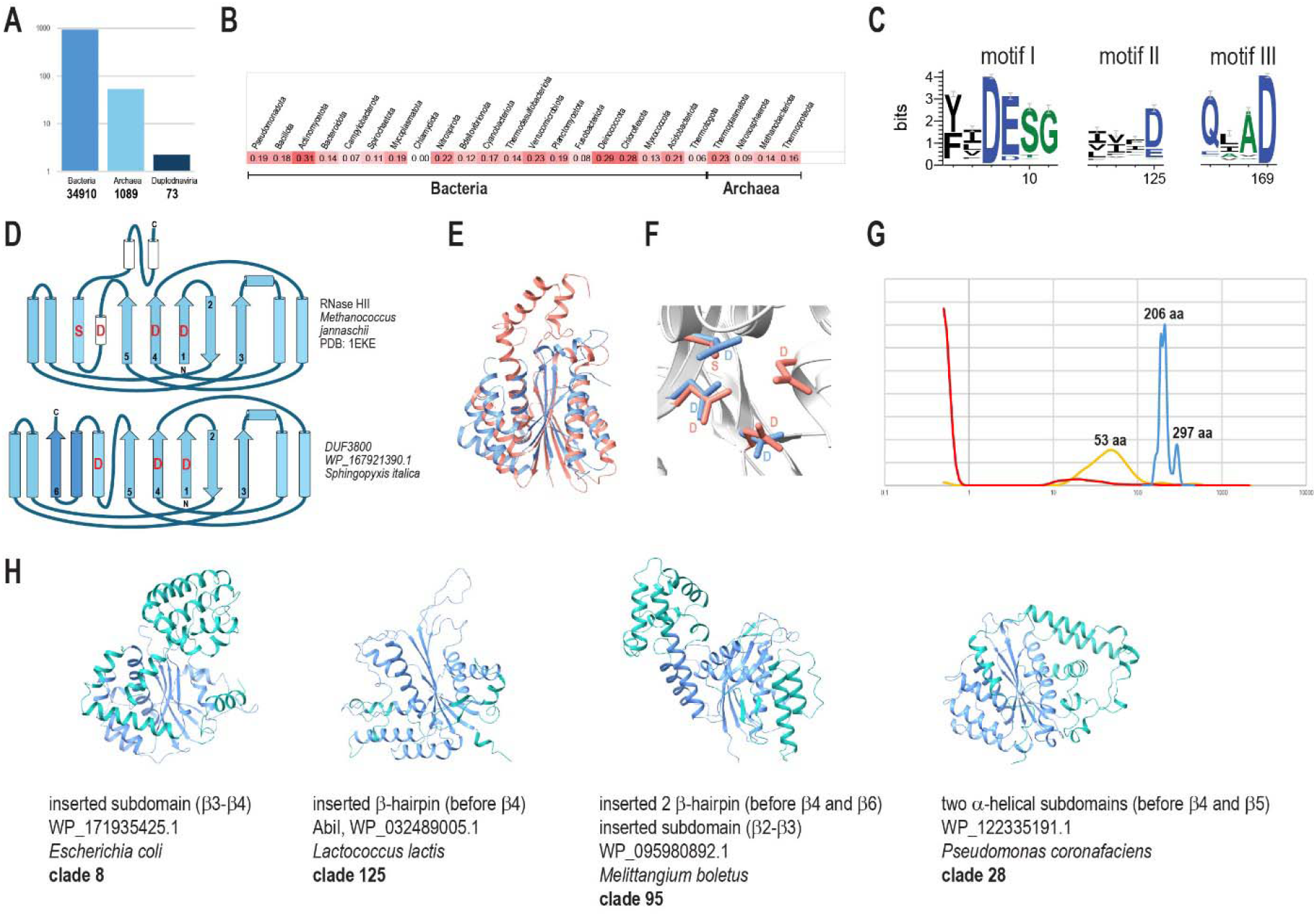
Abundance, sequence features and structure comparison of DUF3800 protein family. (A) The number of identified DUF3800 proteins in cNR database in bacteria, archaea and viruses; Y-axis, fraction of the DUF3800 proteins (promille). The actual counts are indicated below bars. (B) Heat map of the relative abundances of DUF3800 in completely sequenced genomes across major archaeal and bacterial phyla. The heat map shows the weighted fraction (scaled from 0 to 1.0) of DUF3800 presence. For each occurrence of a DUF3800 in a genome, the weight of the respective genome was assigned (see details in Methods). The sum of the weights of the DUF3800 loci was normalized to the sum of the weights across the taxon. (C) Seqlogo for three most highly conserved sequence motifs in DUF3800. The logo was calculated with default parameters using Weblogo server ^106^ for alignment of 500 consensus sequences of core DUF3800 domain built for 500 deep clades in the tree (see also Figure S1); (D) Topology diagram comparing RNase H and DUF3800 core folds. Alpha helices are shown as cylinders and beta-strands as arrows. The strands are numbered in the order from the N-terminus to the C-terminus. The key conserved amino acid residues are shown in red. Light blue denotes shared structural elements, and darker blue denotes elements specific for DUF3800; (E) Superposition of the minimal DUF3800 protein (containing only DUF3800 domain) and RNase HII (pdb: 1EKE); (F) Same superposition as in (E) zoomed in to the catalytic site. (G) Probability density function for the sizes of core DUF3800 domain footprints (blue), N-terminal extension (red) and C-terminal extension (yellow). Typical domain sizes at the peaks are indicated. The X-axes is in a log scale. (H) Examples of various intradomain insertions (cyan) into the core DUF3800 domain (blue). Extra N-terminal elements are also shown in cyan.

### Sequence analysis and structural features of the DUF3800 domain

Multiple alignment of the core DUF3800 domain sequences encompassed several conserved motifs, three of which correspond to the RNase H superfamily catalytic triad (Figure 1C, Figure S1). The consensus motif “**D**ETG…**D/E**…QLA**D**” is similar to that identified in the Pfam database (PF12686) ^42^. In some DUF3800 subfamilies, the second aspartate is replaced by a glutamate and a few subfamilies appear to be inactivated because they lack identifiable conserved negatively charged residues (Figure S1).

Comparison of the AlphaFold structural model for a representative protein (WP_167921390.1 from *Sphingopyxis italica*) containing only the DUF3800 core domain to the PDB database of protein structures yielded a strong match to RNase HII of *Methanococcus jannaschii* (PDB 1EKE, z-score 9.48), with the three conserved aspartates forming a likely catalytic triad (Figure 1D-F). However, the third catalytic aspartate of DUF3800 is located in a different alpha helix than in the RNases H and replaces the serine that is conserved in RNase HI and HII families ^43^. The C-terminal elements are also different in the two structures. In DUF3800, an additional 6^th^ beta strand extends the beta sheet core. This strand is preceded by an alpha helix also belonging to the DUF3800 fold core.

Most of the identified DUF3800 proteins contain only the DUF3800 core domain, with minimal N-terminal and C-terminal extensions, but there is considerable diversity of C-terminal fusions (Figure 1G, Table S1). The size of the DUF3800 core domain itself varies widely, between 123 and 439 aa, with two pronounced peaks at 206 and 297 aa (Figure 1G). The proteins comprising the second peak contain a variety of insertions into the core domain, many of which are located between β3 and β4 strands and are predicted to form an alpha-helical subdomain (Figure 1H).

We reconstructed a phylogenetic tree of 36,094 DUF3800 core domain sequences and used it as a framework for further sequence and neighborhood analysis (Dataset 4). The tree contained 500 deep clades (Table S1). Footprints of multiple alignments of DUF3800 sequences from each of these clades are provided for use as query profiles to search sequence databases for DUF3800 domains (Dataset 1).

We modeled the structures of representatives of all 240 clades that consisted of 10 or more sequences. Comparison of these modeled structures showed that, despite substantial variation, they all formed a connected graph, linked by structural similarity, indicating conservation of the common core fold (TM-scores >0.5 ^44^, Figure S2, Dataset 5). This observation implies that all varieties of DUF3800 descended from the same ancestral core domain, diverging via the acquisition of various intra-domain insertions.

### ncRNAs associated with DUF3800 genes

Eleven distinct non-coding RNAs have been previously identified next to DUF3800 genes, mostly downstream, and were designated DUF3800-I to DUF3800-XI ^36^. One plausible hypothesis is that these RNAs function as antitoxins for DUF3800 toxin RNases in type III TA modules. We scanned the upstream and downstream intergenic regions around DUF3800 genes for homologs of RNAs available in the Rfam database ^45^ and mapped the identified ncRNAs onto the DUF3800 tree (Figure 2A, Figure S3, Table S1). We found that only 7.8% of the DUF3800 genes were associated with ncRNAs assignable to one of the known RNA families. Most of these ncRNAs are encoded downstream of the DUF3800 genes, but DUF3800-VI and DUF3800-XI are typically encoded upstream, and DUF3800-VI RNA is found in upstream and downstream regions equally frequently, and occasionally, in both (Table S1 and Table S3 RNA).

**Figure 2.**
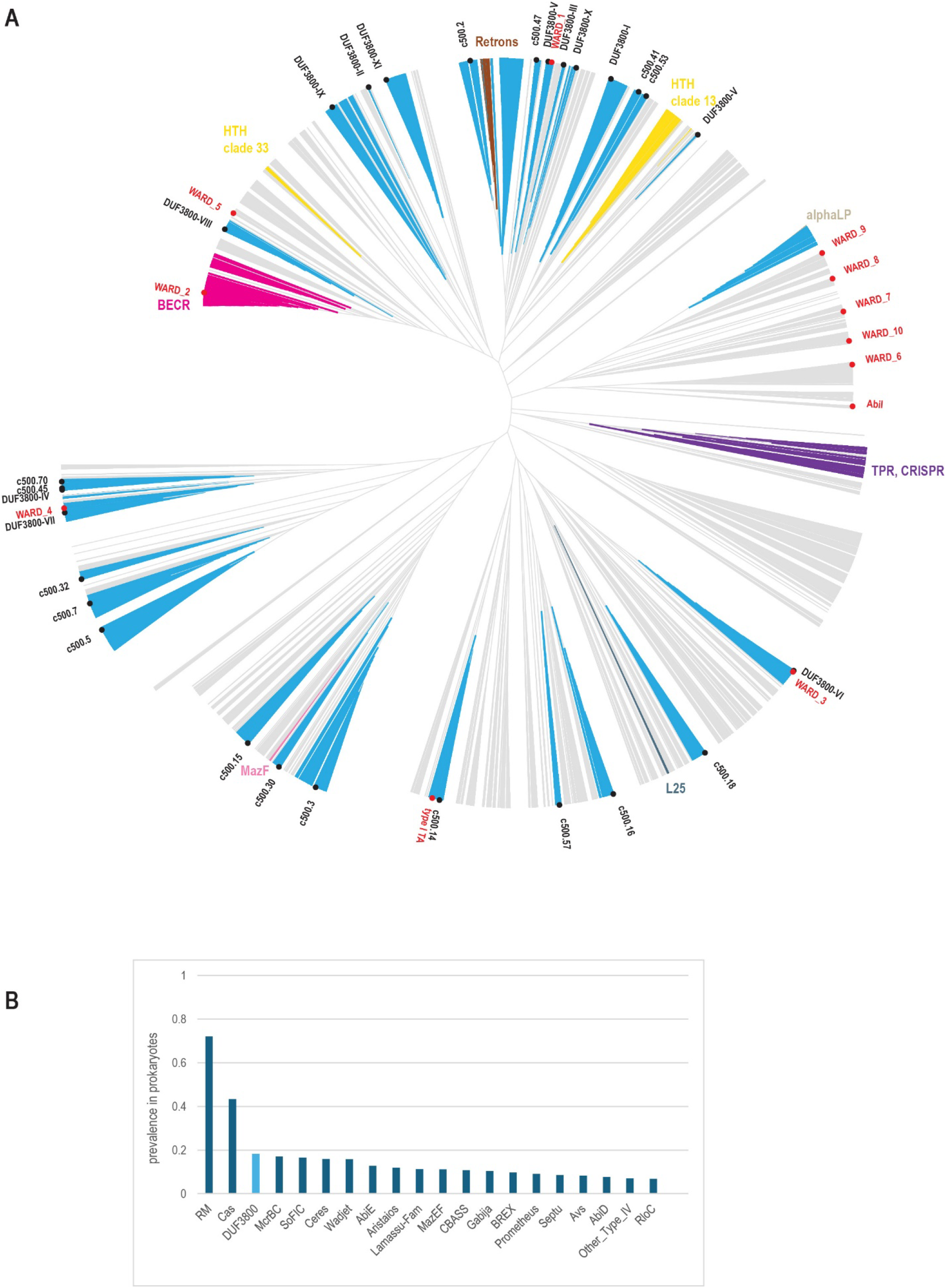
Comprehensive phylogenomic analysis of the DUF3800 family. (A) Phylogenetic analysis of 36,095 DUF3800 cluster representatives (90% clusters in cNR database) using FastTree (see Methods). The tree is unrooted and is depicted as ultrametric. The original tree with branch lengths and bootstrap estimates is available in Dataset 4. Blue indicates 27 clades with more than 100 sequences for which an ncRNA was predicted in at least 30% of the loci. These clades are denoted by either DUF3800 I-XI RNAs or by clade number if potential ncRNAs in these clades were identified in this work (see details in Table S1 and S3). Several major clades not associated with ncRNA are color-coded other than blue; a short clade name is indicated next to each of these clades. Experimentally characterized WARD proteins are indicated by red circles and denoted by a protein name from the literature or a WARD number as shown in the Table S1. (B) Comparison of DUF3800 prevalence (fraction of genomes, corrected for database sampling bias, across the set of 47,545 completely sequenced prokaryotic genomes) with those of known defense systems, all types of RM and CRISPR-Cas systems were combined. Defense systems were identified using DefenseFinder.

Despite the absence of identifiable members of known ncRNA families, many DUF3800 genes are surrounded by long intergenic regions, providing enough room to encode a ncRNA. Considering that most DUF3800 RNAs are shorter than 95 nt and are encoded immediately adjacent to the DUF3800 genes, we calculated the median up- and downstream intergenic region lengths for all 500 deep clades in the tree and used the 95 nt cutoff to identify clades that have enough room for encoding an ncRNA. This analysis suggests that more than 90% of the DUF3800 genes can potentially encode an ncRNA, providing an upper bound estimate (Figure S3 and Table S1). To assess the evolutionary conservation of these regions, which might be indicative of the presence of an RNA gene, we clustered the upstream and downstream regions using a two-step procedure (see Materials and Methods for details) and identified 287 clusters including at least 5 non-redundant sequences each (Dataset 6). Among these clusters, 37 matched 9 of the 11 previously reported DUF3800-associated ncRNAs, whereas 6 were discarded because they either were very short or matched other known RNAs irrelevant in the defense context, such as tRNAs (Table S3 RNA). In the final set of 281 conserved intergenic regions, 139 alignments could be folded into complex RNA structures with the free energy estimate (ΔG) below -20 kcal/mol per 100 nt, and 245 had ΔG equal or lower to known DUF3800 RNAs (Table S3 RNA). These 281 alignments were scanned against the DUF3800 intergenic regions (see Methods), yielding additional 7,455 matches likely representing novel ncRNAs. Altogether, potential ncRNAs could be identified in 29% of the DUF3800 loci and typically are clade-specific (Figure 2A, Figure S3 and Table S1). However, given the presence of long intergenic regions in most of the loci, this is most likely to be an underestimate (Figure S3 and Table S1).

### Abundance, domain architectures and genomic neighborhoods of DUF3800 proteins

We used DefenseFinder ^46^ to identify known defense genes in the DUF3800 neighborhoods (10 genes upstream and downstream) and found that 41% of the DUF3800 genes are located within potential defense islands (Figure S3, Table S1, Dataset 7; see Materials and Methods). This association is not confined to specific DUF3800 clades, but rather spread across the entire tree, suggesting that most of these proteins are involved in defense against mobile genetic elements (MGE). There are, however, several notable exceptions which we discuss below.

Comparison of the abundance of DUF3800 with those of other known defense systems in completely sequenced genomes showed that, by the phylogenetically weighted prevalence, DUF3800 ranked third behind the most common Restriction-Modification (RM), and CRISPR-Cas systems (Figure 2B, Figure S3). Thus, if most of the DUF3800 genes are involved in defense, the largely uncharacterized defense systems that, presumably, employ these nucleases as effectors could be among the most prevalent in the prokaryotic immune repertoire.

Next, we examined domain fusions and putative operons which include DUF3800 (Figure 2A and 3, Figure S3, Table S1, Dataset 7). At least two relatively large DUF3800 subfamilies encompassed fusions with DNA-binding domains (Figure 2A; highlighted in yellow). One of the largest clades in the tree (c500.13) consists of proteins in which DUF3800 is fused to a winged helix-turn-helix (wHTH) domain, most similar to the D3 domain of type IIS restriction endonuclease FokI (Figure S4 and S5). The D3 domain is part of the DNA recognition module of this nuclease, but its specific role is not entirely clear ^47^. Only a few of these DUF3800 proteins are associated with predicted ncRNAs and are not parts of any conserved predicted operons although they are typically encoded within defense islands (Table S1). Recognition of MGE DNA by the wHTH domain might directly activate DUF3800, bypassing the RNA-mediated step.

**Figure 3.**
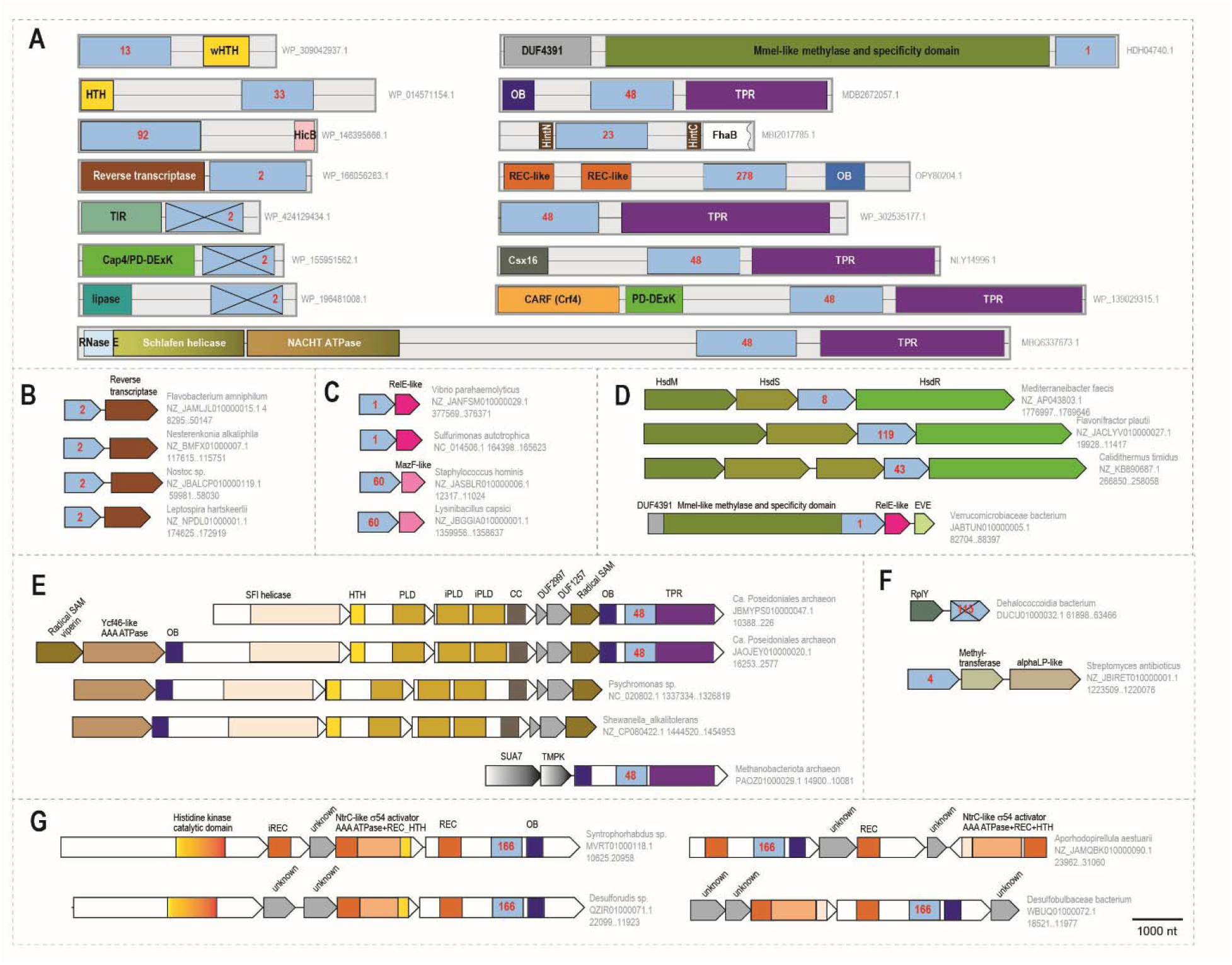
Multidomain proteins and conserved neighborhoods of DUF3800. (A) DUF3800 domains are shown in blue; the clade number to which each domain belongs is indicated inside the box (see Table S1 for clade assignment). Domains are shown roughly to scale and represented by colored boxes. Homologous domains are shown by the same color, and a short description is provided inside the boxes. Protein accessions are indicated on the right; B) Retron-associated neighborhoods (see designations common for panels B-G below); C) Association of DUF3800 with BECR and MazF RNases; D) DUF3800 as a potential component of RM systems; E) DUF3800 as a component of a distinct, complex defense system; F) potential recruitment of DUF3800 for non-defense functions; G) DUF3800 as a component of a predicted signal transduction system. The neighborhoods are shown to the same scale (see scale bar in panel F). For each neighborhood, the species name, nucleotide accession and coordinates are provided. Genes are shown as block arrows. Domains are shown in selected proteins by colored boxes. Homologous genes or domains are shown by the same color. Unknown domains or genes predicted to encode components of the respective systems are shown in gray. Non-defense genes are shown by a gray gradient. Gene names or short descriptions are given above the shapes where available. Abbreviations: OB – OB fold, TPR - tetratricopeptide repeats, HTH – helix turn helix domain, wHTH – winged helix turn helix domain, TIR - Toll/interleukin-1 receptor/resistance domain, PD-DExK – endonuclease of PD-DExK family, REC – phosphoacceptor receiver domain, iREC – inactivated REC domain, PLD – nuclease of phospholipase D family, iPLD – inactivated PLD nuclease; CC – coiled-coil region.

The next most common DUF3800 fusion is with another HTH domain that belongs to a subclade in clade c500.33 and is specific for *Lactobacillaceae*. This domain does not show similarity to any functionally characterized HTH domains. Structural modeling reveals an additional helical bundle subdomain, connected by apparently unstructured loops (Figure S5). Most of these genes are associated with predicted ncRNAs in both upstream and downstream intergenic regions. This HTH domain might function as a self-repressor, as in many TA systems ^20^, and the additional helical subdomain might function as the sensor, whereas the ncRNA could be the antitoxin.

Another notable fusion of DUF3800 is with the HicB domain (Figure S4). These fusion proteins comprise a small subfamily specific to *Planctomycetota*. A DALI search ^48^ with a structural model of one of such proteins produced two significant matches, one to the uncharacterized DUF3234 protein (PDB: 2z0r; TTHA0547), and the other, to HicB (PDB: 6u0i; Figure S5). HicB is the antitoxin of the well-studied type II toxin-antitoxin system HicAB, where HicA is an RNAse toxin ^49^. HicB is an RNAse H-like domain, albeit partial and inactive ^50^. No ncRNA was predicted in these loci. The HicB-DUF3800 fusion appears to be analogous to other toxin-antitoxin fusions of type II TA system components ^51^. Such fusions have not been extensively characterized functionally, and they cannot operate on the same principle as canonical type II TA, which rely on constant resupply of the faster-decaying antitoxin, so they probably represent a yet unknown class of TA systems.

A substantial fraction of DUF3800 genes is associated with retrons, a major group of defense systems that are composed of a reverse transcriptase (RT), a noncoding RNA, and diverse effector proteins ^32,52,53^. This link has been noticed previously, and the DUF3800-associated retrons have been classified as type VII, with subtype A1 encompassing RT fused to DUF3800 and subtype A2, in which DUF3800 is encoded in a separate gene in the same operon with RT ^32^. In agreement with previous observations, the retron-associated DUF3800 domains form a distinct clade in our tree, with two subclades corresponding to the A1 and A2 subtypes (Figure 2A, Figure 3A and 3B, Table S1). In these systems, DUF3800 likely functions as the effector activated by interaction with the nucleic acid synthesized by the RT in response to infection. In addition, we identified a small branch within the retron clade A2 where DUF3800 is likely inactivated, but remarkably, is fused to 6 other, distinct potential effector domains, namely, NAD-depleting TIR domain, Cap4-like PD-DExK superfamily DNAse, putative lipase of alpha-beta hydrolase family, CHAT family cysteine protease, trypsin-like serine protease and a phosphorylase (Figure 3A, Table S1). These domains function as effectors in a variety of unrelated prokaryotic defense systems including CRISPR-Cas, NLR and others ^54–58^. In these strikingly complex defense systems, the inactivated DUF3800 domain likely plays a structural role in the organization of the effector complex and/or signal transduction.

The largest clade in the tree (c500.1) corresponds to the putative two-gene operons encoding a BECR (Barnase, EndoU, Colicin E5/D, and RelE)-fold nuclease, in addition to DUF3800 (Figure 3C). RelE, an interferase that cleaves mRNAs inside the ribosome, is one of the most abundant toxins in type II TA systems ^20,59^. The BECR-fold proteins in the DUF3800 loci encompass a distinct pattern of conserved putative catalytic residues, different from that in the known RelE toxins (Table S1, Figure S4 and S5). Their closest relative is the poorly characterized protein YjaZ that, in *E. coli*, has been shown to affect growth in the stationary phase via an unknown mechanism ^60^. The sequence of YjaZ matches the pattern of putative catalytic residues in DUF3800-associated BECR proteins, suggesting that these proteins are active RNases, functionally linked to the DUF3800 RNase.

In addition to the DUF3800-BECR connection, we identified a smaller clade where DUF3800 is encoded in a putative operon with a MazF-like protein (Table S1). MazF is another well-studied RNase toxin from type II TA systems ^20,59^. MazF-like proteins encoded in DUF3800 loci are predicted to be active because they retain the two catalytic residues shown to be essential for the RNase activity ^61^ (Figure S4 and S5). *In silico* co-folding of both protein pairs, DUF3800-BECR and DUF3800-MazF, did not produce stable complexes, suggesting that these proteins do not strongly interact strongly. Considering the absence of identifiable ncRNA in the DUF3800-BECR and DUF3800-MazF loci, it seems likely that one of the RNases is the antitoxin that might specifically cleave the toxin mRNA, as shown for the only functionally studied type V TA system ^62^. In that system, the toxin GhoT is a membrane peptide, whereas the antitoxin GhoS is an RNAse that cleaves the GhoT mRNA. So far, however, no TA systems consisting of two RNases have been identified.

It has been previously observed that many genomic loci encoding immunity systems also encompass genes that induce cell dormancy or death ^6^. One of the explanations of this phenomenon is that toxins can be activated to arrest virus production and buy time for immunity systems, such as RM or CRISPR-Cas, to clear the virus. Some evidence supporting this hypothesis has been recently obtained, namely, it has been shown that TAs function in concert with RM systems ^63^. Another, perhaps, more common scenario is that the TA is the second line of defense that is activated and causes dormancy or cell death when the active defense fails, in particular, being blocked by a viral antidefense protein ^6,8^. Our analysis of DUF3800 loci showed that RM systems are the most common neighbors of DUF3800 (Table S1 and Table S4), and the latter are often embedded into otherwise typical type I RM operons (Figure 3D). In some cases, this arrangement is conserved in minor clades (e.g. c500.43, Table S1), suggesting co-evolution of RM and DUF3800. Typically, for such operons, we could not predict a ncRNA, and the intergenic regions are short, suggesting that DUF3800 is co-expressed with the RM system and its toxicity can be harnessed to provide time for the RM system to act or to trigger abortive infection when RM fails. In another, even more intriguing case, DUF3800 is linked to an RM system component in a small branch within the BECR-DUF3800 clade (c500.1, Table S1). This variant comprises a predicted operon consisting of three genes encoding a multidomain protein containing DUF3800, RM-like DNA methylase and DUF4391 domain, a BECR nuclease, and occasionally, a protein containing an EVE domain, which is typically involved in modified DNA recognition ^64,65^ (Figure 3D). The methylase and its cognate specificity domain are most similar to those of the type IIL RM MmeI, but the characteristic N-terminal DNase domain of MmeI ^66^ is replaced by the DUF4391 domain that is not similar to any known nucleases. This variant appears to be a derivative of the minimal DUF3800-BECR system, where one RNase (likely, DUF3800) is the toxin and the other the antitoxin, possibly, degrading the toxin mRNA. The site-specific DNA methylase might add a regulatory or signaling dimension to the DUF3800 toxicity, especially, in conjunction with the affinity of the EVE domain for modified DNA bases.

Another small but notable branch in the tree corresponds to the fusions of DUF3800 with OB fold domains and TPR repeats (Figure 3A, Table S1). Most of the OB fold domains bind nucleic acids, whereas TPR repeats are involved in protein-protein interactions and are common sensor domains in prokaryotic Avs (Antiviral STAND ATPases) ^57,67^. In several uncultured archaea of phylum *Thermoplasmatota*, this protein is a component of a highly complex predicted defense system containing, additionally, a radical SAM enzyme, an AAA ATPase of the Ycf46 family, a superfamily I helicase, two proteins containing PLD family nuclease domains (one of these with two inactivated PLD domains), and two uncharacterized proteins ^68^ (Figure 3A and 3E, Figure S4). Helicases, radical SAM enzymes and PLD nucleases are components of other known defense systems, whereas an ATPase of Ycf46 family has been shown to contribute to the response to various environmental stresses ^69–75^. In several archaeal genomes, the DUF3800-TPR-OB protein is encoded as a singleton, and the remaining part of the system is present in some bacteria that lack this fusion protein, so the link between the two components of this system seems to be specific for *Thermoplasmatota* (Figure 3E). Given the presence of the TPR repeat module that likely functions as a sensor of viral proteins, this system might be functionally analogous to the Avs.

Some of the DUF3800 genes are not obviously linked to defense. A small *Chloroflexota*-specific clade (c500.115) includes inactivated DUF3800 proteins that are encoded in a putative operon with ribosomal protein L25, the only ribosomal protein that is upregulated under stress, suggesting that the inactivated DUF3800 is also involved in stress response ^76^ (Figure 3F). A larger clade represented in many actinomycete genomes consists of DUF3800 associated with an alpha-lytic protease and a methyltransferase (Figure 3F, Table S1). The protease belongs to a well-studied family of serine proteases ^77^, whereas the methyltransferase showed no close relationships with functionally characterized ones, but was predicted to be active (Figure S4). In this clade, several ncRNAs have been predicted in the DUF3800 intergenic regions (Table S1). Another system is highly complex and, based on gene content, is apparently involved in signal transduction (Figure 3G). In this case, the DUF3800 domain is part of a large, multidomain protein also containing OB-like and phosphoacceptor receiver (REC) domains (Figure S5). The locus additionally contains genes encoding a highly diverged histidine kinase fused to a PAS domain, an NtrC-like σ54 activator AAA ATPase, which is fused to a REC and HTH domains, a protein with an inactivated REC domain and various uncharacterized protein, sharing little or no sequence similarity with each other. Histidine kinases and REC domains are typical components of two-component signal transduction systems ^78^. Generally, a sensor domain fused to a histidine kinase recognizes a signal and activates the kinase which phosphorylates the REC domain, which, in turn, activates an effector. In this case, one of the effectors appears to be DUF3800 that most likely cleaves RNA. The second potential effector is related to a σ54 activator AAA ATPase, which is predicted to activate transcription of the targeted genes ^79,80^. Given that the classic σ54 is involved in stress response ^81^, this system is likely to perform a similar function, but the predicted RNA cleavage by DUF3800 is a distinct feature that remains to be investigated.

Another case of probable function switch is the replacement of a homing endonucleases of group I introns, typically of LAGLIDADG family ^82^, by DUF3800 (MBI2017785.1, Figure 3A). There are at least 5 such caes, found only in MAGs (metagenomically assembled genomes), including several *Parcubacteria* from the Candidate Phyla Radiation (CPR) superphylum ^83^. In addition, inteins containing the DUF3800 domain are inserted into an adhesin-like exoprotein (e.g. MBI2017785.1) or chitinase (TSC94215.1) (Table S1). It remains to be determined if the proteins of this DUF3800 subfamily cleave dsDNA and function as homing endonucleases.

Should this be the case, these DUF3800 nucleases could expand the armory of nucleases employed for targeted genome modifications ^84^.

### DUF3800-TPR fusions and the link to CRISPR-Cas systems

One of the major clades in the tree includes mostly proteins with the DUF3800 domain fused to TPR (tetratricopeptide repeats) solenoid domains (Figure 2A; highlighted in purple). TPR domains are well-characterized alpha helical repeat modules involved in protein-protein interaction and molecular pattern recognition ^85^. Notably, TPR solenoids are often fused to STAND ATPase domains and are responsible for recognition of conserved viral proteins by the Avs ^57^. We found that 47% of the DUF3800-TPR proteins in our dataset are encoded in the vicinity of CRISPR-Cas systems or CRISPR arrays (Figure 4A and 4B), often in the same operon with *cas* and CRISPR-Cas ancillary genes and, in a few cases, fused to Cas proteins (Figure 3A and 4B, Table S1, Dataset 7). This clade also includes two branches consisting of proteins not fused to TPR, but the larger of these branches retains association with CRISPR-Cas systems (Figure 4A). The sequences and domain architectures of the DUF3800-TPR proteins in this clade show considerable diversity, and many contain an additional, N-terminal helical domain which often is fused to a diverged CARF (CRISPR-associated Rossmann fold)-like domain (Figure 4C, Table S1, Figure S3). Most often, genes encoding DUF3800-TPR proteins are associated with type III systems or their components, especially, CARF-domain proteins, which are key sensors in cOA (cyclic oligoadenylate) signaling by CRISPR-Cas type III systems (Figure 4A and 4B) ^58,86^. In this signaling pathway, Cas10 synthesizes cOA upon recognition of foreign DNA or RNA, and the CARF domain binds this messenger, activating an effector, typically, a HEPN RNAse ^58^. The activated HEPN domain cleaves RNA non-specifically, causing growth arrest, whereas a ring nuclease provides negative feedback by cleaving cOA and eventually shutting down the response ^33,86,87^. The function(s) of the DUF3800-TPR proteins in type III CRISPR-Cas systems is unclear, but notably, some of these proteins (eg. MFY9179919.1) are fused to a Csx16-like domain, a ring nuclease ^88^ (Figure 3A, Table S1). Thus, at least in some cases, the DUF3800 domain might function as an additional effector, with the nuclease activated by a signaling molecule produced by Cas10, and the ring nuclease attenuating the cell damage by inactivating the signal ^89^.

**Figure 4.**
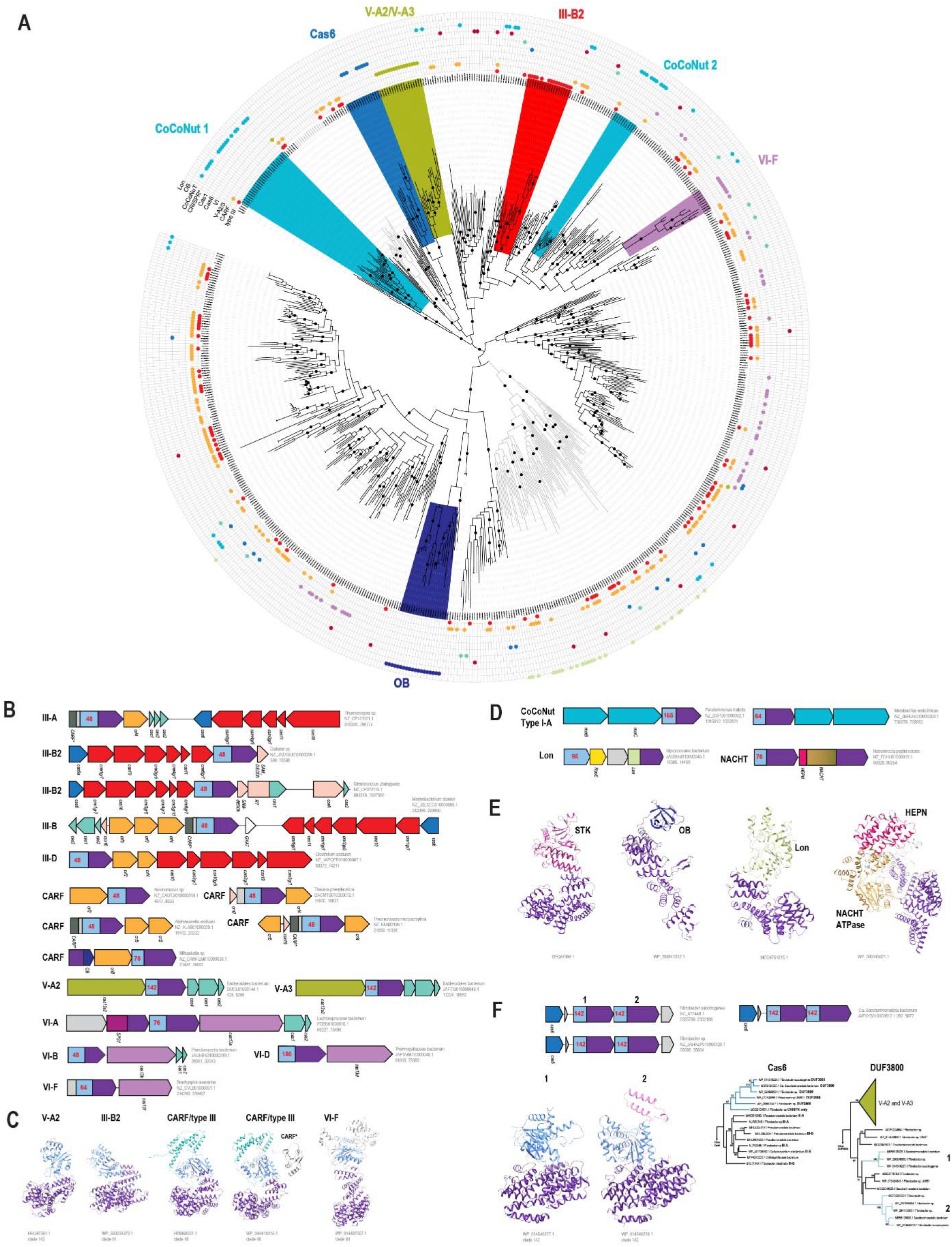
DUF3800 proteins containing TPR domains. (A) A subtree corresponding to the clade of the DUF3800 domain tree (Figure 2A) which is enriched in proteins fused to TPR domains. Branches consisting of DUF3800 domains fused to TPR domains are shown in black, and branches consisting of proteins without TPR domains are shown in grey. Bootstrap support values greater than 75% are indicated by black dots on the respective branches. Clades are colored and a short description is provided for each. The colored dots in the circles around the tree indicate the presence of the genes or defense systems in the respective DUF3800 loci, according to the labels indicated for each circle. CRISPR* denotes the presence of a CRISPR array only or other *cas* or ancillary genes not shown in other circles (for details see Table S1 and Dataset 7). B) DUF3800 domains associated with CRISPR-Cas systems and their ancillary genes. The designations are the same as in Figure 3, but the genes that belong to multigene systems, such as type III effector complex and CRISPR spacer adaptation module, are all shown by the same color, and description of each gene is provided below the arrows. The *cas* genes are named according to the published profiles ^33^, gr5 and gr7 mean Cas5 and Cas7 components of the class 1 effector complexes; C) AF3 models for DUF3800-TPR proteins associated with CRISPR-Cas systems. The alpha-helical domain often found in type III/CARF loci is shown by cyan; CARF* is a diverged variant of the CARF domain; D) Non-CRISPR conserved neighborhoods; E) AF3 models of TPR-containing proteins fused to other effectors but often encoded in the vicinity of DUF3800 genes; F) Cas6-associated DUF3800-containing system contains two DUF3800 genes denoted as 1 and 2. The operon organizations, structural models of the two DUF3800 paralogs (pink – insertion into the DUF3800 fold, which structurally distinguishes the two DUF3800 paralogs) and subtrees for Cas6 and DUF3800 with closest relatives are shown. For DUF3800, the subtree corresponds to the respective branches in the main tree (Figure 2A, Table S1, Dataset 4). The branches corresponding to DUF3800 1 and DUF3800 2 from the respective loci are shown by teal and light blue, respectively. The sister branch for DUF3800 associated with V-A2 and V-A3 is collapsed and colored moss green. Cas6 phylogeny was built using FasTree program for closest Cas6 homologs identified in the cNR database. The clade corresponding to Cas6 associated with the DUF3800 is colored blue. The genes or CRISPR-Cas system types found in the vicinity of the *cas6* genes are indicated by bold.

Association of DUF3800-TPR genes with type VI CRISPR-Cas systems is the second most frequent (Figure 4A and 4B). The effector of the type VI systems is Cas13, a protein containing two HEPN domains which targets RNA ^90^ and, more specifically, induces cell damage via tRNA cleavage ^91^. Also, as mentioned above, DUF3800-TPR genes are associated with type V-A2 CRISPR-Cas systems that cause cell damage via collateral RNA and ssDNA cleavage upon target DNA recognition ^92,93^. Recently, a type V-A3 variant associated with DUF3800-TPR has been discovered, where the effector protein, Cas12a3, cleaves tRNA tails upon recognition of the target RNA ^94^. Notably, all CRISPR-Cas systems found to encompass DUF3800-TPR target RNA, compatible with the possibility that DUF3800 RNase is an additional effector, likely, causing cell damage.

In the phylogenetic tree of DUF3800, the CRISPR-associated DUF3800-TPR proteins form three strongly supported clades, suggesting co-evolution of specific DUF3800 subfamilies with the corresponding CRISPR-Cas variants (Figure 4A,B). Of special note is the DUF3800-TPR subfamily associated with the III-B2 variant. In the III-B2 CRISPR-Cas systems, Cas10 produces SAM-AMP that functions as a signaling molecule instead of cOA ^95^. Typically, III-B2 systems lack CARF domains and employ different effectors that sense SAM-AMP, such as membrane protein CorA ^95^. Thus, this particular DUF3800-TPR protein seems to be a strong candidate for the role of another SAM-AMP-sensing effector. Most of the other DUF3800-TPR proteins seem to be promiscuous and are likely activated by RNA cleavage products of different CRISPR-Cas systems. Indeed, the first insights into activation of DUF3800-TPR proteins have been recently reported ^96,97^. It has been shown that the TPR domain of a DUF3800 protein linked to a V-A3 CRISPR-Cas system binds tRNA CCA-tail, a product of tRNA cleavage by Cas12a3, and activates DUF3800 ^96^. The activated DUF3800 nuclease cleaves cellular mRNAs and causes cell dormancy, complementing tRNA cleavage by Cas12a3 and providing efficient protection against phages. It seems likely that DUF3800-TPR nucleases in other type III systems are activated in a similar manner because HEPN domains fused to CARF also can cleave tRNA ^98^. Given that tRNAs have a universal CCA tail, activation of DUF3800 is expected to be highly robust regardless of which defense system produced these molecules.

CRISPR-Cas are not the only defense systems linked to DUF3800-TPR genes. The DUF3800-TPR tree includes two clades where these proteins are encoded in putative operons with CoCoNuT I-A systems (Figure 4B). The CoCoNuT systems were discovered by genome analysis and have not yet been experimentally characterized ^99^. They are apparently derived from well-characterized type IV RM systems (McrBC) which recognize and cleave modified viral DNA. Type I-A is a minimal CoCoNuT system and, based on the presence of a PD-DExK nuclease domain, can be predicted to target DNA. However, many McrB-like components (CnuB) of type I and other CoCoNuT systems contain an OB-fold domain suggesting that these proteins bind RNA. Thus, the CoCoNut-associated DUF3800-TPR proteins associated with these systems might also sense the same RNA molecules.

There are also a few more associations we identified for the proteins that belong to the clade shown in the Figure 4A. One corresponds to OB-DUF3800-TPR proteins associated with a complex system described above (Figure 3E). In another branch, the DUF3800 proteins lack TPR but, in addition to various CRISPR systems or their known ancillary proteins, are often encoded in the vicinity of a Lon-like serine protease fused to a TPR domain (Figure 4D and 4E). It seems likely that, in this case, the TPR domain performs the same function as in DUF3800-TPR proteins. Lon-TPR and DUF3800-TPR are not the only TPR-containing proteins found in extended DUF3800 loci from this clade. Serine-threonine kinases encoded in some type IV-A CRISPR-Cas loci also contain TPRs (Figure 4B and 4E) one of which has been shown to sense tRNA tails ^97^. TPR domains are also present in NACHT NTPases (a family in the STAND superfamily) and, in a few occasions, a DUF3800 domain is fused to NACHT and Schlafen domains (Figure 4D and 4E). In eukaryotes, Schlafen is an important component of translation control under stress conditions that cleaves tRNA and rRNA via a RNAse E-like domain ^100^.

Recently, Schlafen proteins have been characterized in bacteria as well, where they are fused to a variety of effector and sensor domains and protect bacteria from viruses through tRNA cleavage by the RNase E-like domain ^101^. This RNase E-like domain is conserved in several DUF3800 fusions, so that such proteins contain two distinct RNase domains (Figure 3A, Figure S4). Some of the TPR domain-containing proteins, such as TPR-OB domain fusions, have been previously identified as significantly associated with type III CRISPR-Cas systems before ^102^. Generally, unlike the TPR domains in Avs that recognize structural patterns in viral proteins, many if not most of the numerous TPR domains linked to various effectors, and DUF3800 in particular, likely sense RNA molecules, in particular, tRNA tails ^96,97^. Additionally, DUF3800 operons related to the non-TPR clade often encode sigma subunit of RNA polymerase RpoE, a stress response associated sigma factor ^103^. RpoE is expected to be involved in regulation of expression of these proteins (Figure 4C).

Finally, we identified a small clade corresponding to several loci from *Fibrobacterota* which encode a Cas6 protein and two DUF3800-TPR proteins (Figure 4F). The two DUF3800-TPR are diverged but apparently originated from a duplication and contain closely similar TPR repeats; the only notable difference between the two paralogs is the presence of a helical insertion in one of the DUF3800 domains (Figure 4F). These DUF3800 proteins form a sister group to a CRISPR-Cas V-A-associated clade which, together with the Cas6 association, suggests that the DUF3800 domain ancestral to these two clades was already linked to CRISPR-Cas systems (Figure 4F). The function of Cas6 in CRISPR-Cas systems is to cleave pre-crRNA, producing mature crRNA which, when bound to effector complexes of class 1 CRISPR-Cas systems, serves as the guide recognizing matching protospacers in foreign DNA or RNA ^104^. The Cas6 protein associated with DUF3800 is most closely similar to type III Cas6 from which it likely evolved (Figure 4F). However, the DUF3800-Cas6 loci do not contain detectable CRISPR repeats or conserved intergenic regions. It seems likely that Cas6 associated with DUF3800 also cleaves RNA, with the cleavage products sensed by the DUF3800-TPR proteins, activating the RNase activity of DUF3800. However, the nature of the Cas6 substrate in this case remains unclear.

### DUF8203 family

DUF8203 family is a diverged version of DUF3800 (HHpred probability 76% for WP_167921390.1 as a query) which is found almost exclusively in archaeal genomes, from both thermophiles (eg. Thermococci) and mesophiles (eg. Halobacteria) (Table S2). Altogether, we identified 152 members of this family in cNR. DUF8203 shares the common core RNase H fold with DUF3800 and similar motifs, including three conserved aspartates (Figure S6). In addition, DUF8203 proteins contain a C-terminal alpha/beta domain (Figure S6). We could not identify homologs of any of the known DUF3800-associated ncRNA next to the DUF8203 genes and detected only three upstream or downstream regions conserved in five or more genomes. However, nearly all DUF8203 genes are flanked by long upstream and/or downstream regions (median 235 nt and 254 nt, respectively), so there is room for encoding an ncRNA (Table S2). Of the 152 identified DUF8203 genes, only 13 are encoded in the vicinity of known defense systems although it should be taken into account that archaeal defense systems remain far less thoroughly explored than bacterial ones. No other conserved neighborhoods were identified either. Nevertheless, given the conservation of the RNase H fold with intact catalytic motifs and the patchy distribution in archaeal genomes, it appears most likely that DUF8203 is a TA module and/or is involved in anti-MGE defense.

## Experimental validation of the defense function for DUF3800 proteins

For an initial experimental characterization of the DUF3800 family, we focused on the simplest variants of putative DUF3800 defense systems. To this end, we selected one representative from each of 10 deep clades, three of which were associated with ncRNAs, one was linked to putative BECR RNase, and all others were standalone genes (Figure 5A). These genes stood out because the majority of DUF3800 proteins are encoded by genes that belong to predicted operons and are inferred to function as effectors of multicomponent defense systems, such as retrons and CRISPR-Cas, suggesting novel modes of defense. Furthermore, the selected DUF3800 genes are not located within typical defense islands, but instead, are associated with phage genes, often appearing to reside within prophages. Defense genes are known to be enriched within prophages, and previous analysis demonstrated the presence of DUF3800 in the P2 but not the P4 defense hotspot ^105^. Seven of the 10 selected proteins were from *E. coli* strains but shared, at most, 19% amino acid sequence identity, highlighting the high level of divergence within the DUF3800-family and suggestive of independent acquisition of these genes via HGT from different sources.

**Figure 5.**
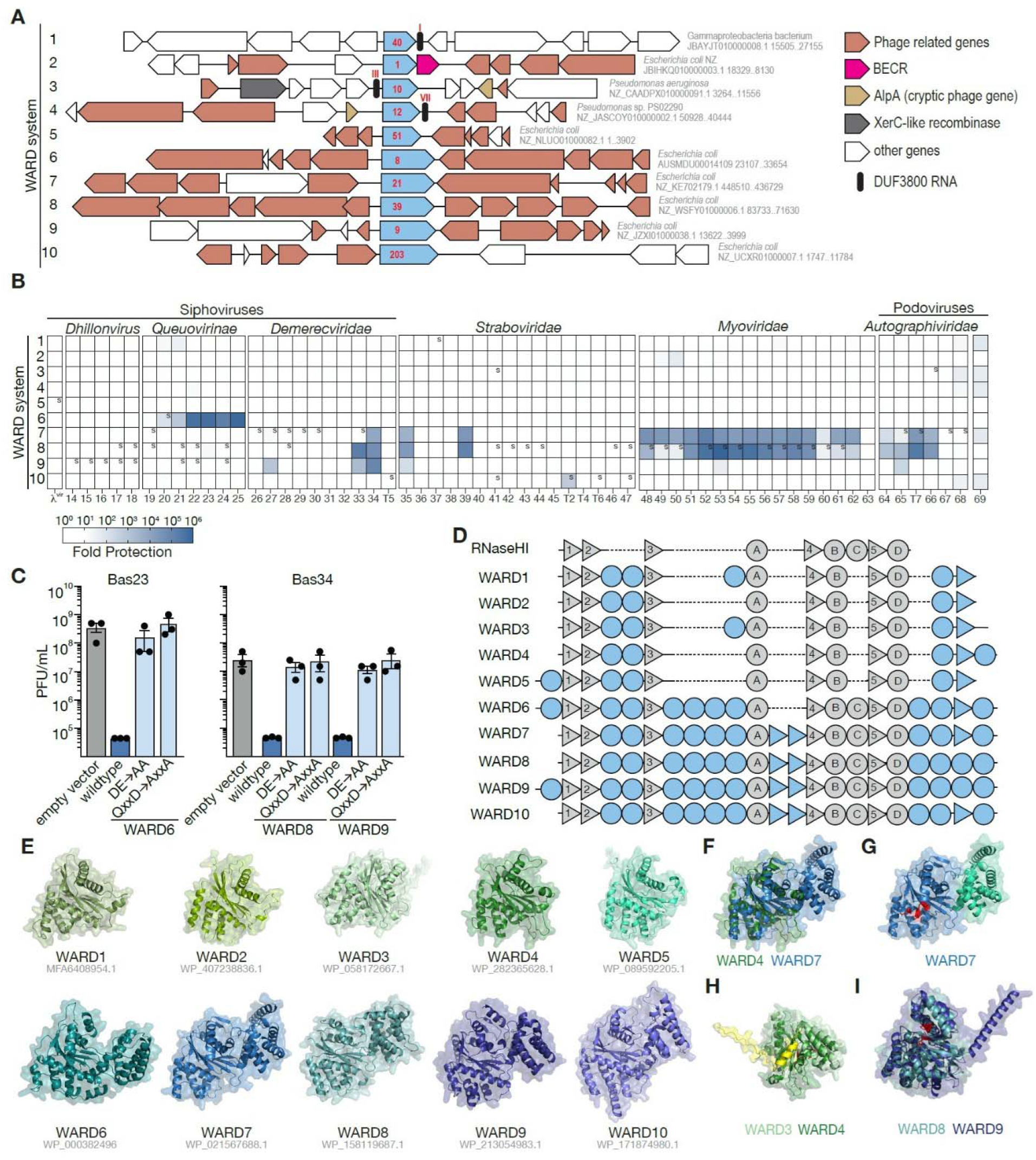
WARDs as single protein defense systems that depend on the nuclease activity of DUF3800 for antiphage defense. (A) Graphical representation of the 10 tested predicted defense systems along with their genomic contexts depicted roughly to scale (see Table S5 for accession numbers and surrounding protein annotations). The DUF3800 domain containing proteins are shown in blue, and the clade number to which the given domain belongs is indicated inside the box in red (see Table S1 for clade assignment). DUF3800 RNAs are shown in black. Phage-related genes are shown in peach; AlpA proteins are shown in light brown; BECR (Barnase, EndoU, Colicin E5/D, and RelE)-fold nucleases are shown in pink; XerC-like recombinases are shown in grey, and other proteins in white. For each WARD, the species name, nucleotide accession and coordinates are provided. (B) Fold-protection (reduction in phage titer) conferred by the indicated WARD system, expressed from the plasmid pLOCO3 in *E. coli* MG1655, against a panel of 62 phages. Phages are ordered by relatedness with phages from the BASEL collection denoted by the Bas#. A small *s* in a cell indicates an observed plaque size reduction in at least two of the biological replicates. The data in the heatmap represent the mean of three independent biological replicates. (C) Resulting plaque forming units (PFU) per mL of the indicated phage when infecting *E. coli* expressing different WARDs with the indicated genotype. The indicated values represent the mean +/- the standard error of the measurement. (D) Cartoon depiction of the secondary structures (as predicted by AlphaFold3) of the WARD proteins highlighting the conserved RNase H-like core fold in grey and unique domains in blue. Triangles represent beta sheets and circles represent alpha helices. RNaseHI; WP_000917883.1. (E) AlphaFold3 prediction of each WARD protein; accession numbers are indicated. All proteins are similarly aligned, centered at the core RNaseH-like fold. (F) Overlay of WARD4 and WARD7 predicted structures highlighting the globular insert domain unique to WARD6-10. Proteins are colored as in E. (G) AlphaFold3 prediction of WARD7 with the DE and QxxD catalytic residues shown in red and the domain unique to WARD6-10 proteins shown in cyan. (H) Overlay of WARD3 and WARD4 with the DE and QxxD catalytic residues shown in red and the unique C-terminal extension colored yellow. Proteins are colored as in E. (I) Overlay of WARD8 and WARD9 with the DE and QxxD catalytic residues in red highlighting the unique N-terminal alpha helix found in WARD9. Otherwise, proteins are colored as in E.

We expressed each of these genes in *E. coli* MG1655 under their native promoters and challenged the resulting strain with a panel of 62 diverse phages. Consistent with the predicted defense function of DUF3800, all but one of these genes provided defense (defined as > 1 log fold protection) against at least one phage (Figure 5B). We therefore propose to rename the DUF3800 domain family <u>W</u>eaponized <u>A</u>ntiphage <u>R</u>Nase-dependent <u>D</u>efense (WARD) family.

Four of the WARDs, WARD6-9, conferred particularly robust defense (up to a million-fold protection), with WARD6 specifically targeting *Queueovirine*, whereas WARDs 7, 8, and 9 targeted numerous unrelated phages (Figure 5B). Notably, WARD6-10 all belong to the same major branch in the tree, together with AbiI (Figure 2A). WARD10 specifically defended against T2 but none of the closely related *Straboviridae.* WARD2, which is in an operon with a BECR domain-containing protein showed mild protection against Bas50. Mutation of the predicted catalytic residues that define the DUF3800-family in WARDs 6, 8 and 9 (Figure 1C) resulted in loss of defense, consistent with the hypothesis that the defense function of DUF3800 proteins depends on their RNase activity (Figure 5C-D).

Notably, the DUF3800 genes that conferred the strongest protection against phages were not associated with ncRNAs and encoded substantially longer proteins (368-406aa) compared to those that did not confer defense or only restricted phage plaque size (224-267aa) (Figure 5A-B). Examination of AlphaFold models showed that, whereas all 10 tested WARDs shared a conserved RNase-H like core fold composed of five beta-strands surrounded by six alpha helices, the larger proteins possessed an additional globular domain inserted between strands 3 and 4 of the core domain and composed of six predicted alpha-helices (Figure 2H, Figure 5D-F, Figure S4). This additional domain was not significantly similar any known domains and is not in contact with the WARD active site (Figure 5G). Comparison of the predicted structures of the diverse WARDs revealed additional features unique to specific systems. WARD3 and WARD4 contain an extended C-terminus compared to WARD1,2 and 5 (Figure 5D-E). In WARD4, this extension is predicted to adopt a helical structure whereas in WARD3 it appears to be unstructured (Figure 5H). WARD5,6 and 9 all possess an N-terminal alpha-helix absent in other WARDs (Figure 5D), which is particularly pronounced in WARD9 (Figure 5I). All these additional domains are not predicted to interface with the active site pocket (Figure 5G-I), suggesting they are involved in sensing phage infection or mediating functionally relevant protein-protein interactions independent of the enzymatic activity.

Previous work has shown that DUF3800 can become toxic and cause cell growth arrest dependent on it’s RNase activty^34,35,96^. Together with the organization of many WARD loci resembling that of type I and type III TA systems, these findings suggest that WARDs mediated defense via abortive infection. Surprisingly, however, liquid infection assays showed that WARDs had minimal impact on bacterial growth, while effectively limiting phage replication, even up to the multiplicity of infection (MOI) of 10 (Figure 6A,D). To further validate this finding, we monitored the amount of phage produced over the first four hours of infection. Whereas phage replication in control cells was readily detectable, replication in WARD-expressing *E. coli* was completely blocked and, on the contrary, a consistent decrease in phage levels was observed (Figure 6B,E). Propidium iodide (PI) staining confirmed that bacteria expressing WARDs remained viable following infection (Figure 6C,F) whereas PI concentration clearly increased in control cells. However, at MOI of 10, we observed a minor but clear impact on the bacterial viability, suggesting that more than direct phage targeting was at play (Figure 6A,C,D,F). Future studies are needed to determine the exact mechanism underlying WARD-mediated defense and at what stages of phage replication WARD acts. In summary, our analysis suggests that DUF3800-containing proteins are commonly involved in antiphage defense via their RNase activity.

**Figure 6.**
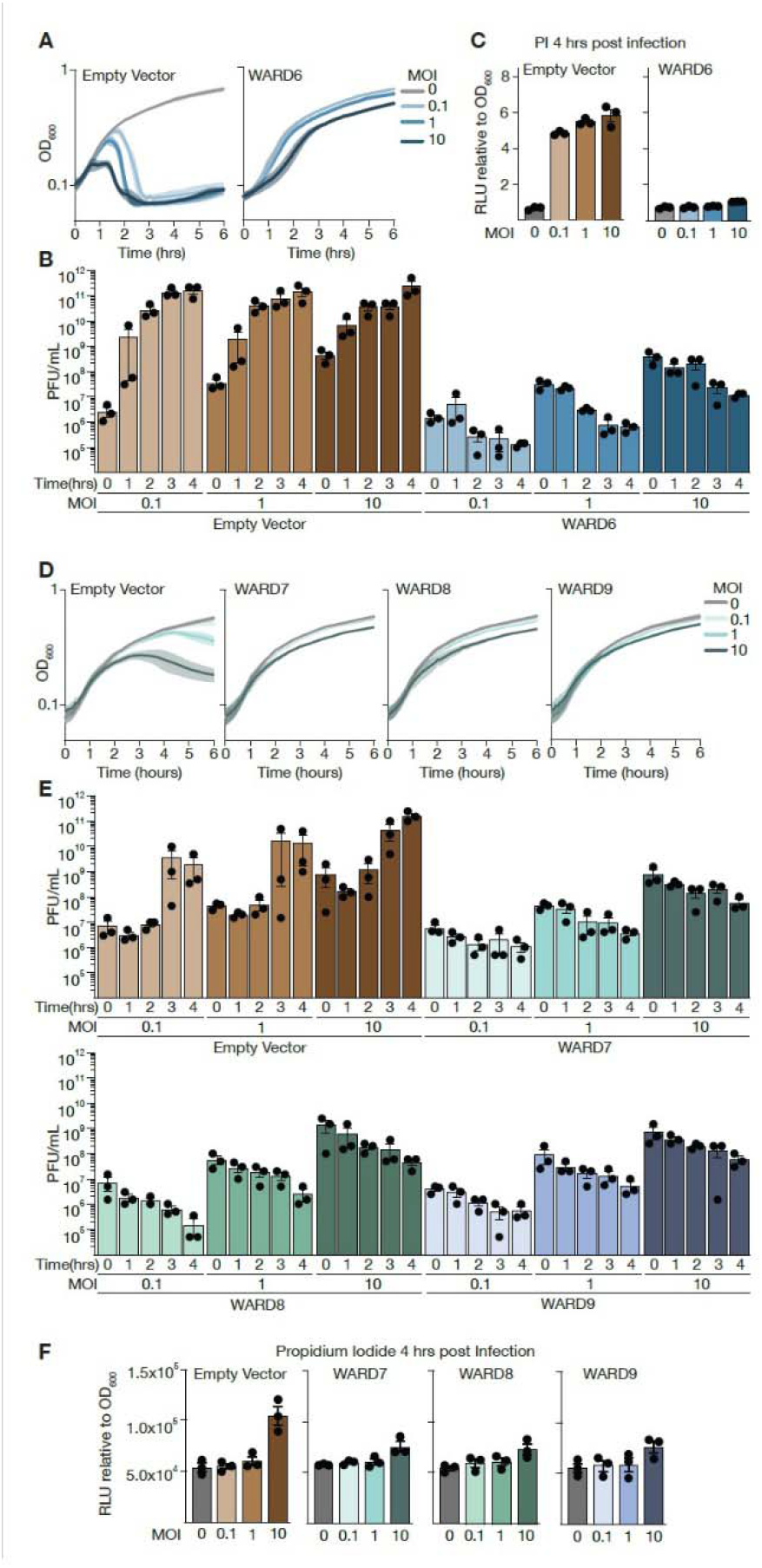
WARDs protect *E. coli* against phages without significantly impacting host bacterial growth. (A) Growth curves of *E. coli* MG1655 expressing sfGFP (empty vector) control (left) or WARD6 (right) upon infection with phage Bas23 at increasing multiplicity of infections (MOIs), measured as optical density (OD_600_) over time. Curves show the mean and standard deviation from three independent biological replicates. (B) Resulting plaque forming units (PFU) of Bas23 recovered from the infection of the indicated host strains over the first four hours of infection at varying MOIs. Data are the mean +/- the SEM. (C) Propidium iodide levels in bacteria expressing the sfGFP or WARD6 and infected with Bas23 at the indicated MOI four hours post infection. Fluorescence is normalized by the OD600 measures. Data are the mean +/- the SEM.(D) Growth curves of *E. coli* MG1655 expressing sfGFP (empty vector) control (left) or the indicated WARD (right) upon infection with Bas34 at increasing MOIs, measured as OD_600_ over time. Curves show the mean and standard deviation from three independent biological replicates. (E) Resulting plaque forming units (PFU) of Bas34 recovered from the infection of the indicated host strains over the first four hours of infection at varying MOIs. Data are the mean +/- the SEM. (F) Propidium iodide levels of bacteria expressing sfGFP or the indicated WARD and infected with Bas34 at the indicated MOI four hours post infection. Fluorescence is normalized to OD600 measures. Data are the mean +/- the SEM.

## Conclusions

In this work, we describe a vast family of bacterial and archaeal proteins that contain a distinct variant of the RNase H fold and are encoded in about 15% of the available complete prokaryotic genomes. Most of the members of this family are encoded in defense islands, many in close proximity of known anti-MGE defense systems, hence the designation WARD (<u>W</u>eaponized <u>A</u>ntiphage <u>R</u>Nase-dependent <u>D</u>efense). One of the WARD proteins has been shown to act as a toxin in a type I toxin-antitoxin module, and an association between a small group of WARDs and defensive retrons has been noticed previously, but otherwise, despite its high abundance, the WARD family has been so far overlooked, apparently, due to the low sequence conservation. Many of the genes encoding WARDs are linked to (putative) ncRNAs suggesting they are toxins of type I or type III TA systems. Unusually for TA loci, however, the ncRNA, a putative antitoxin, is, in most cases, encoded downstream of the WARDs. The most common genomic context of the WARDs is a predicted two-gene operon, where the second gene encodes another RNase of the BECR or MazF families. These two-component systems might be type V TA modules in which one of the RNase, most likely, WARD, is the toxin, and the other one is the antitoxin, possibly, cleaving the toxin mRNA.

Proteins containing WARD domains display a remarkable variety of domain architectures, some containing large inserts, primarily, with predicted alpha-helical structure, within the core RNase H fold, and others fused to additional domains. A widespread WARD subfamily features the combination of WARD with a C-terminal TPR domain. The WARD-TPR proteins are encoded within the same genomic loci with a variety of known defense systems, most often, type III, V-A and VI CRISPR-Cas. The WARD-TPR proteins might function as additional effectors in these systems, with the TPR domains possibly activating the WARD RNase upon sensing RNA cleavage products.

We experimentally validated the defensive capacity of 9 of the 10 tested standalone WARDs from different *E. coli* strains against a broad variety of phages and demonstrated the dependence of the anti-phage protection on the catalytic residues of the RNase H fold. Nine out of 10 tested WARs are likely encoded within prophages (Table S5). The WARDs with the most robust antiphage activity belong to a distinct clade in the tree together with AbiI (Figure 2A).

Perhaps, unexpectedly, these WARDs were not associated with ncRNA but rather contained a large alpha-helical insert within the core RNase H fold. These additional domains might function as infection sensors activating the WARD RNase. None of the tested WARDs caused measurable cell damage in *E. coli*, indicating that, unlike most known TA modules, these are immune systems directly clearing phage infection.

In addition to the WARDs proper, we identified AWARDs (Archaeal WARDs, DUF2803), distant relatives of the WARD family encoded in a variety of archaeal genomes and predicted to protect these organisms against archaeal viruses.

After hundreds of distinct defense systems have been discovered in bacteria and archaea during the last few years ^4^, it appeared that the widespread immune mechanisms were already known, with only rare defense systems remaining to be discovered. However, the WARDs represent a striking exception, apparently, being the third most abundant type of defense systems in prokaryotes, after RM and CRISPR-Cas. The diversity of the domain architectures of the WARD proteins and the genomic contexts of WARD genes imply a broad variety of anti-MGE defense mechanisms that remain to be explored. More generally, the discovery of the WARDs suggests that widespread but highly divergent defense systems might still be lurking in prokaryotic genomes.

## RESOURCE AVAILABILITY

### LEAD CONTACT

Further information and requests for resources should be directed to and will be fulfilled by the lead contacts, Eugene V. Koonin, Kira S. Makarova and Hannah E. Ledvina.

### MATERIALS AVAILABILITY

Plasmids, bacterial strains, and phages used in this study are listed in Tables S6 and are available upon reasonable request.

### DATA AND CODE AVAILABILITY

All sequence data analyzed in this study were obtained from NCBI GenBank (downloaded May 2026). All data supporting the findings of this study are available within the article and its supplemental information. Dataset are deposited on the following FTP site: https://ftp.ncbi.nih.gov/pub/wolf/_suppl/duf3800/

Custom scripts used for data analysis are available from the authors upon reasonable request.

### AUTHORS’ CONTRIBUTIONS

Conceptualization, K.S.M., Y.I.W, H.E.L. and E.V.K.; investigation, K.S.M., Y.I.W, K.A, P.V.J., K.K.A, K.A.K, and R.Q.T.; methodology, K.S.M., Y.I.W, H.E.L.; data curation, K.S.M., Y.I.W, K.A., H.E.L.; writing—original draft, K.S.M., Y.I.W, H.E.L., E.V.K.; writing—review and editing, K.A., K.K.A, K.A.K, R.Q.T.; supervision, project administration, and funding acquisition, K.S.M., E.V.K. and H.E.L.

## ACKNOWLEDGMENTS

The authors thank the Aaron Whiteley lab for their support and thoughtful discussion of this project. Y.I.W, E.V.K. and K.S.M were supported by the Intramural Research Program of the National Institutes of Health (NIH). The contributions of the NIH authors are considered Works of the United States Government. The findings and conclusions presented in this paper are those of the author and do not necessarily reflect the views of the NIH or the U.S. Department of Health and Human Services. K.A.K. was supported in part by T32GM149391.

## STAR Methods

### Identification of DUF3800 and DUF8203 homologs

PSI-BLAST ^107^ search with e-value threshold of 0.0001 using the position-specific scoring matrix (PSSM) for DUF3800 (pfam12686) as the query was used to identify an initial set of homologs of DUF3800 in the May 2026 version of NCBI clustered NR database (non-redundant protein sequences clustered with 90% sequence identity and 90% coverage). The set was complemented by results of five separate PSI-BLAST searches initiated from the following queries: AbiI (WP_032489005.1, positions 1-80 aa), Retron-associated DUF3800 (WP_123861595.1, positions 1-80 aa), CRISPR-Cas V-A2-associated DUF3800 (MCI7571711.1, positions 1-80 aa), CRISPR-Cas VI-F-associated DUF3800 (WP_069732041.1, positions 166-266 aa). These PSI-BLAST searches were performed iteratively online for 10 iterations with no composition-based statists correction and inclusion threshold e-value of 0.01 or until first false positives appear with E-value <0.01. The combined protein set was additionally examined for false positives using HHpred search against PDB, PFAM and CDD databases with default parameters; the presence of any DUF3800 hit with no probability value threshold was required to include sequence in the seed set.

Then, several iterations of the following procedure were performed: DUF3800 domain footprints were extracted from the full -length protein sequences and clustered using MMSEQS2 ^108^ with similarity threshold of 0.2 and coverage threshold of 0.75. Sequences within the clusters were aligned using FAMSA ^109^; alignments were examined for the cluster-wide presence of the conserved catalytic motifs and converted into MMSEQS2 profiles and into PSI-BLAST PSSMs. First pass searches against the NCBI clustered NR database were performed using MMSEQS2 at sensitivity 7.5 and default e-value threshold; second pass searches were performed against the MMSEQS2 hits using PSI-BLAST with the respective iteration PSSMs to extract the footprints and e-value threshold of 0.0001. In the end, this procedure produced 36,103 distinct proteins, containing a full-length DUF3800 domain.

DUF8203 homologs were identified using online PSI-BLAST search against the clustered NR database for 5 iterations.

### Multiple alignment and phylogenetic analysis

DUF3800 domain footprints were extracted from the full -length protein sequences and clustered using MMSEQS2 with similarity threshold of 0.2 and coverage threshold of 0.75.

Sequences within the clusters were aligned using FAMSA. The cluster alignments were iteratively merged using the following procedure: first, alignment profiles were compared to each other using HHSEARCH ^110^ in the global alignment mode; pairs with high mutual length coverage (coverage threshold of 0.75) were grouped and progressively aligned to each other using HHALIGN. After each step, alignments were filtered, with positions containing more than 2/3 gap characters removed, reducing the alignment to the common conserved core. These alignments were again compared to each other using HHSEARCH and the process was repeated until all clusters were aligned.

The resulting global MSA was examined for the correct alignment of the N- and C-terminal catalytic motifs (or the corresponding sites in the inactivated protein sequences). Several incorrectly aligned groups were extracted, re-aligned from the footprints and examined using HHSEARCH and HHPRED to identify the (approximate) location of long insertions into the DUF3800 domain core. After removal of insertions, these alignments were re-aligned to the master alignment using HHALIGN.

Approximate maximum-likelihood tree was constructed from the global alignment of the DUF3800 domain using FastTree ^111^ with gamma-distributed site rates and WAG evolutionary model.

### Genomic neighborhoods of DUF3800

36,094 of the 36,103 identified DUF3800-containing proteins from the clustered NR database mapped to at least one source contig in GenBank. The genomic neighborhoods of DUF3800 genes, consisting of up to 10 annotated protein-coding genes upstream and downstream (fewer, if limited by the contig end), were extracted. A minimal sequence set was compiled, containing all 36,094 DUF3800 genes and consisting of 35,784 representative genomic neighborhoods, encoding 559,358 distinct proteins. Among these, 36,094 proteins represented the core set of DUF3800-containing proteins; additional 146 proteins, not represented in clustered NR database, also contained full-length DUF3800 domains and another 130 proteins contained partial DUF3800 domains.

All proteins in these neighborhoods were annotated with CD, COG and Pfam profiles from the NCBI CDD database ^112^ and custom Cas protein profiles ^33^ using PSI-BLAST searches with e-value threshold of 0.0001. The entire neighborhoods were also annotated using DefenseFinder ^46,113^.

Deep clustering of non-DUF3800 proteins was performed in a two-step procedure. First, all these proteins were subject to 3 iterations of MMSEQS2 search against themselves with sensitivity parameter of 7.5, and clustered based on the search results using MMSEQS2 “clust” command. First-level cluster alignments were produced from protein sequences using FAMSA; then, alignments containing sequences that are immediate neighbors of DUF3800 genes were searched against all alignments using HHSEARCH; clusters that produced hits with at least 0.75 mutual length coverage were merged.

Regions covering up to 10 kbp up- and downstream of DUF3800 genes, were searched for CRISPR arrays using MinCED 0.4.2 tool ^114^.

### Protein structure analysis

For selected DUF3800 proteins and products of neighboring genes, the structures were predicted using AlphaFold3 ^115^ and visualized using ChimeraX ^116^. Structural comparison was performed using Foldseek ^117^ and Dali ^48^.

Protein structures for all clusters of DUF3800 sequences that included 10 or more representative proteins (240 of the 500 clusters) were predicted using ESMfold2 ^118,119^. The sequence most similar to the cluster alignment consensus was selected as the master sequence (except for the 10 clusters that included proteins chosen for the experimental validation, where the corresponding WARD1-WARD10 proteins were selected as the master sequences). The 240 predicted structures were compared to each other using FoldSeek ^117^ with alignments used as the input; the estimated TM scores ^44^ were recorded for each pair; the higher of the two values (one normalized by the query length, the other by the target length) was taken to represent the comparison result.

### Prediction of non-coding RNAs in DUF3800 intergenic regions

Intergenic regions up- and downstream of the annotated DUF3800 genes were identified and extracted. Regions shorter than 30 bp were discarded; long sequences were truncated to 500 bp. A search for Rfam covariance models ^45^ was performed in these sequences using Infernal with default parameters ^120^.

Additionally, the extracted intergenic region sequences were clustered using MMSEQS2 with similarity threshold of 0.5 and coverage threshold of 0.75; clusters, containing 5 or more sequences were aligned using MAFFT ^121^, reduced to positions with less than 10% gaps, converted to HMM (Hidden Markov Models) using Infernal and searched all against all using cmscan (HMM search only) with default E-value 0.02 ^122^. All sequences from alignments that passed the threshold were combined, realigned and new HMM models were generated to replace those from the original alignments.

All HMM models were used to scan DUF3800 intergenic regions using cmscan. Consensus sequences from the corresponding alignments were searched against the Rfam covariance models to establish operational equivalence between Rfam profiles and locally generated alignments. Then, the results of the two scans of the DUF3800 intergenic regions (Rfam covariance models and local HMMs) were combined.

Consensus RNA secondary structures for these alignments were predicted using the RNAalifold program of the ViennaRNA package ^122^.

### DUF3800-containing proteins in completely sequenced prokaryotic genomes

DUF3800 footprints were matched to sequences in the database of proteins, encoded in 47,545 completely sequenced prokaryotic genomes, downloaded from the NCBI Genomes FTP site in November 2023, using MMSEQS2 (sensitivity parameter of 1). This search produced 4,787 unique hits, covering more than 75% of the query length, mapped to 10,102 distinct genes in 8,389 genomes.

Relative genome weights ^123^, accounting for the uneven genome sampling across the tree of life, were derived from the global genome phylogeny, based on concatenated alignments of phylogenetic markers ^124^.

For quantitative comparisons, defense systems in completely sequenced genomes were identified using DefenseFinder ^46^.

### Bacterial strains and growth conditions

*E. coli* strains used in this study are listed in Table S6. *E. coli* were grown in LB medium (1% tryptone, 0.5% yeast extract, 0.5% NaCl) shaking at 37 °C, 220 rpm unless otherwise noted. Where applicable, media were supplemented with carbenicillin (100 μg ml−1) to ensure plasmid maintenance. All *E. coli* strains were stored in LB plus 30% glycerol at −70 °C. *E. coli* OmniPir ^125^ was used for plasmid construction and propagation. *E. coli* MG1655 (CGSC6300) ^126^ was utilized for all experimental data.

### Plasmid construction

All plasmids used in this study are listed in Table S6. The coding sequences for candidate defense proteins, along with both upstream and downstream genetic regions were obtained as synthetic DNA and cloned into the SbfI and NotI sites of pLCOO3 ^127^. DNA sequences were cloned into destination vectors using 18–25 bp overhangs and Gibson Assembly. Point mutations were cloned by mutagenic PCR and isothermal assembly. Clones were transformed either into a modified strain of OmniMax *E. coli* (Invitrogen) by electroporation plated on LB with the appropriate selection. Positive clones were verified by whole plasmid sequencing (Plasmidsaurus). Prior to use in downstream phage experiments, sequence-verified plasmids were transformed into MG1655 by electroporation and plated on LB with the appropriate selection.

### Phage amplification and storage

Phages used in the study are listed in Table S6 ^128^. All phage lysates were generated from *E. coli* MG1655 using modified double agar overlay plate amplification as previously described ^129^. Briefly, stationary phase MG1655 was infected with roughly 10,000 plaque-forming units (PFU) of phage in LB + 0.35% agar, 10 mM MgCl_2_, 10 mM CaCl_2_, and 100 μM MnCl_2_. Plates were incubated overnight at 37 °C and the following day phages were collected by adding 5 ml of SM buffer (100 mM NaCl, 8 mM MgSO_4_, 50 mM Tris-HCl pH 7.5, 0.01% gelatin) directly to the plate, incubating for around 30 minutes at room temperature, then collecting and chloroform treating the resulting liquid. Lysate titers were determined by spotting a serial dilution of the phage onto 0.35% LB agar plus 10 mM MgCl_2_, 10 mM CaCl_2_, and 100 μM MnCl_2_ containing stationary phase MG1655. Plates were incubated overnight at 37 °C and the resulting titer in PFU ml−1 was calculated. Phage stocks were stored at 4 °C in SM buffer.

### Phage infection assays

Phage defense activity of the candidate defense proteins was evaluated using a double-agar overlay spot assay as previously described ^125^. Overnight cultures of *E. coli* MG1655 cells expressing the individual candidate proteins were diluted into fresh LB medium with appropriate selection and cultured to mid-logarithmic phase. For each assay, 400 µL of the mid-log culture was mixed with LB + 0.35% agar containing additional 10mM MgCl_2_, 10mM CaCl_2_ and 0.1mM MnCl_2_. This mixture was poured over an LB + 1.5% agar plate to make a uniform bacterial lawn.

Phage stocks were serially diluted 10-fold in SM buffer. 2 µL of each dilution was spotted on the overlay. After the spots had absorbed into the overlay, plates were incubated overnight at 37°C. Plaques were counted the following day.

Phage titers were determined from dilutions containing countable plaques and expressed as plaque-forming units per milliliter (PFU/mL). When a dilution produced a diffuse or hazy zone of clearing rather than clearly resolved individual plaques, the lowest phage concentration that produced visible clearing was assigned a value equivalent to 10 plaques for titer estimation^2^. When no plaques or visible clearing were observed at any dilution, a value of 0.9 plaques at the least diluted spot was used to represent the assay detection limit ^125^.

Log Fold protection was calculated by dividing the PFU of a given phage on the strain expressing the empty vector (pLOCO3-sfGFP) by titers obtained on the strain expressing the candidate defense proteins. All experiments were performed three independent times. Full datasets are provided in Table S7 and S8.

### Phage kill curve and viral titer during infection

Overnight cultures of the indicated bacterial strains were diluted into LB supplemented with 10 mM MgCl_2_, 10 mM CaCl_2_, and 0.1 mM MnCl_2_ to a starting OD_600_ of 0.05. Cultures were grown for an hour and normalized by OD_600_ to 0.1 and distributed into a 96-well plate^4^. Phages were then added in equal volumes resulting in the indicated multiplicity of infection (MOI). Following phage addition, cultures were transferred to a Tecan Spark Cyto plate reader, and OD_600_ was recorded at 5-minute intervals for 6 hours.

To quantify phage replication, samples were collected from infected cultures at hour intervals postinfection for a total of 4 hours and treated with 1:10 volumes of chloroform to kill bacteria ^130^. Samples were serially diluted and spotted onto the phage-sensitive empty vector host using the agar overlay method described above. Full datasets are provided in Table S8.

### Membrane permeability

Membrane permeability was monitored in parallel with the OD_600_ using propidium iodide (PI). PI was added to the cultures at a final concentration of 5 µg/mL ^131^, and fluorescence was recorded every 5 minutes over the course of infection using the same plate reader. For simplicity we have only present to values at 4 hours post infection; however, the complete dataset can be found in Table S8.

## Supplementary Information and Datasets

**Figure S1.**
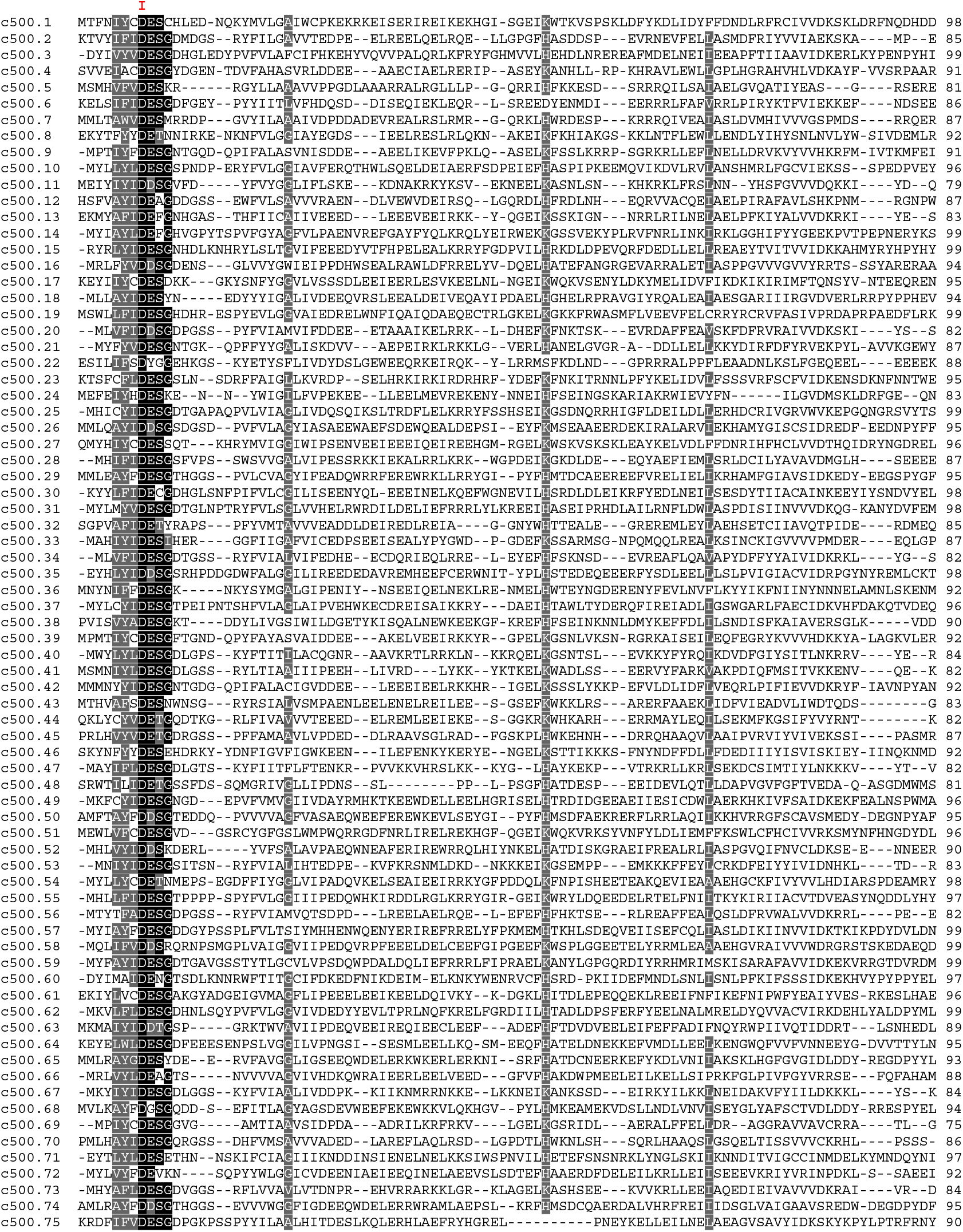
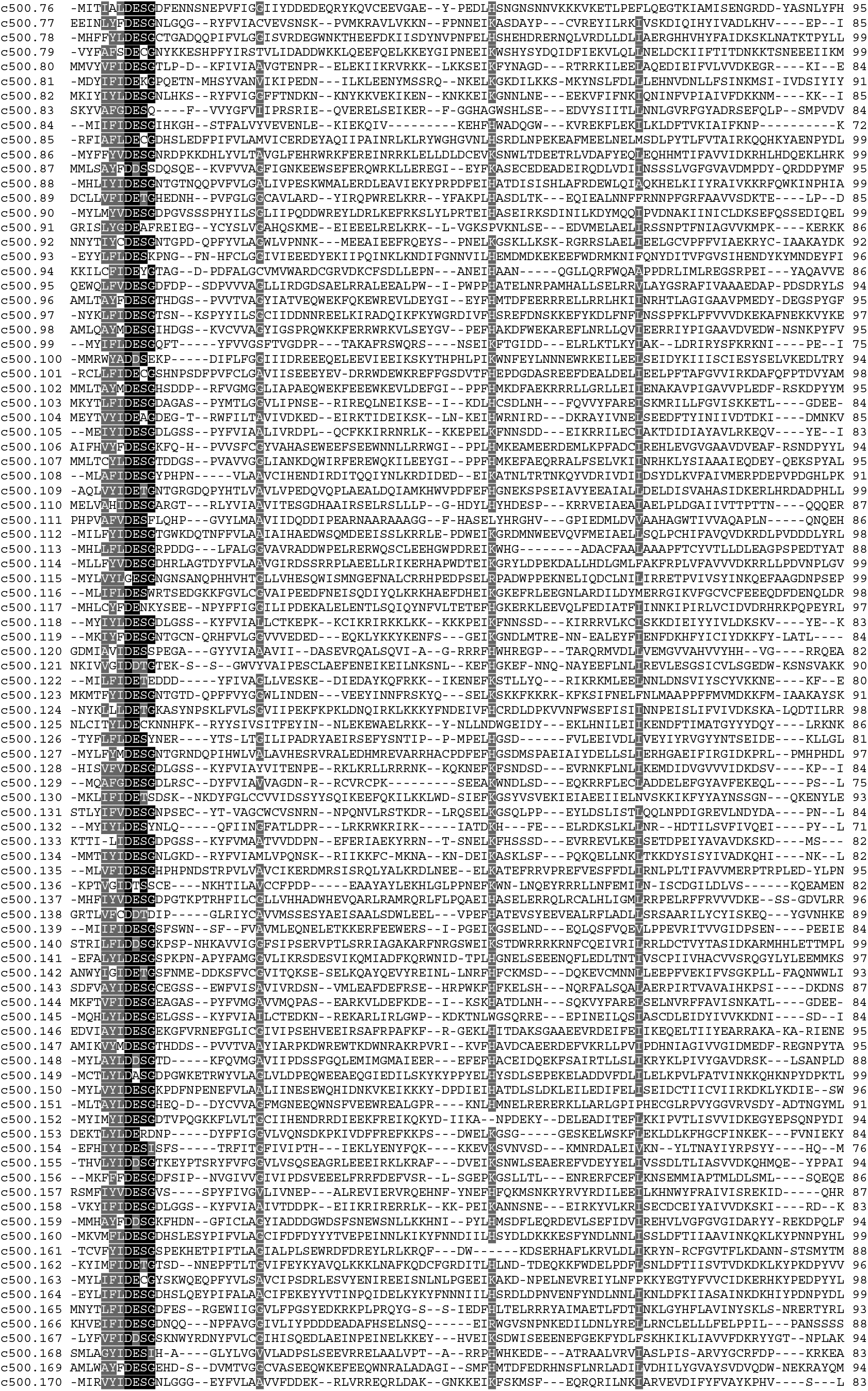
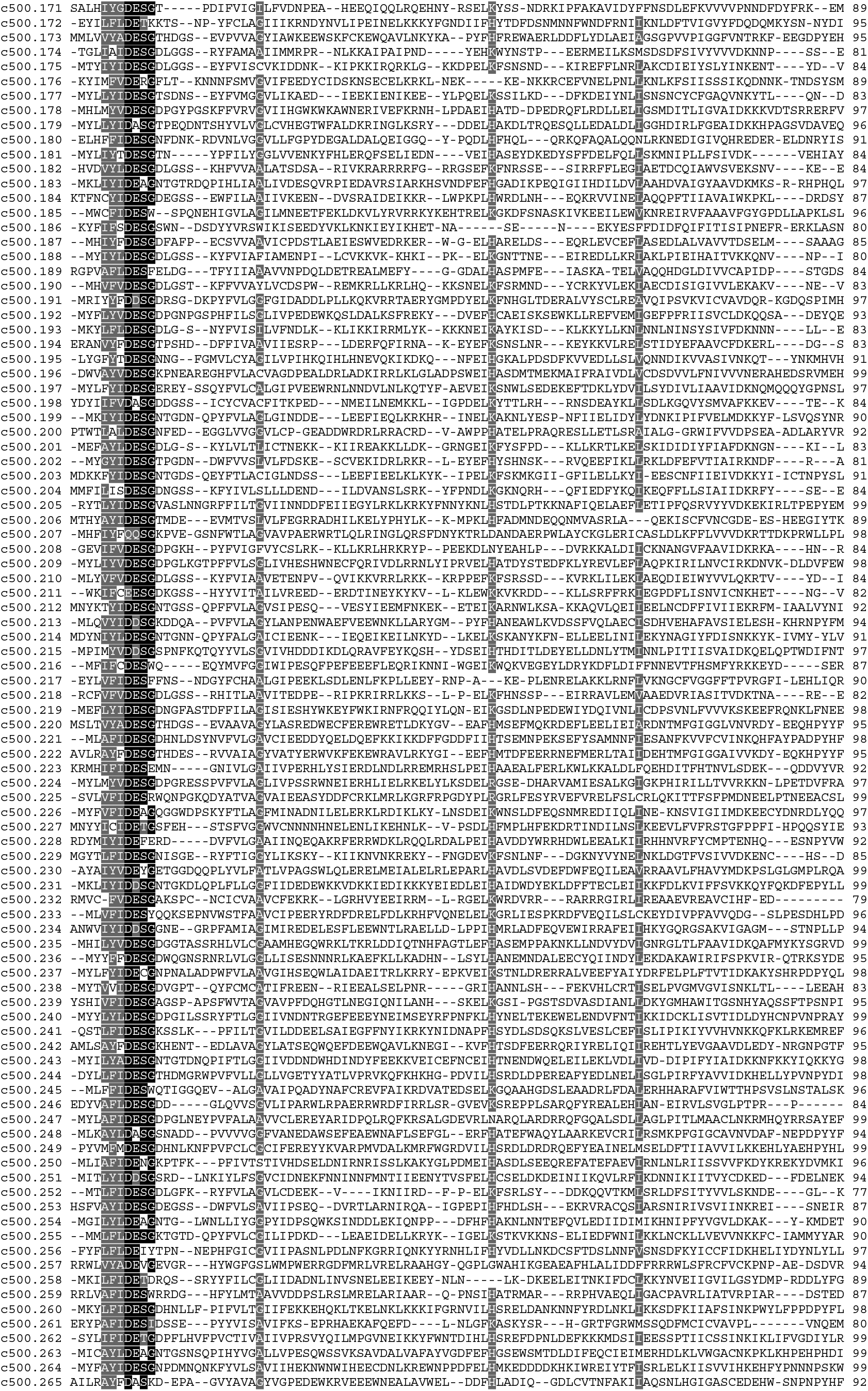
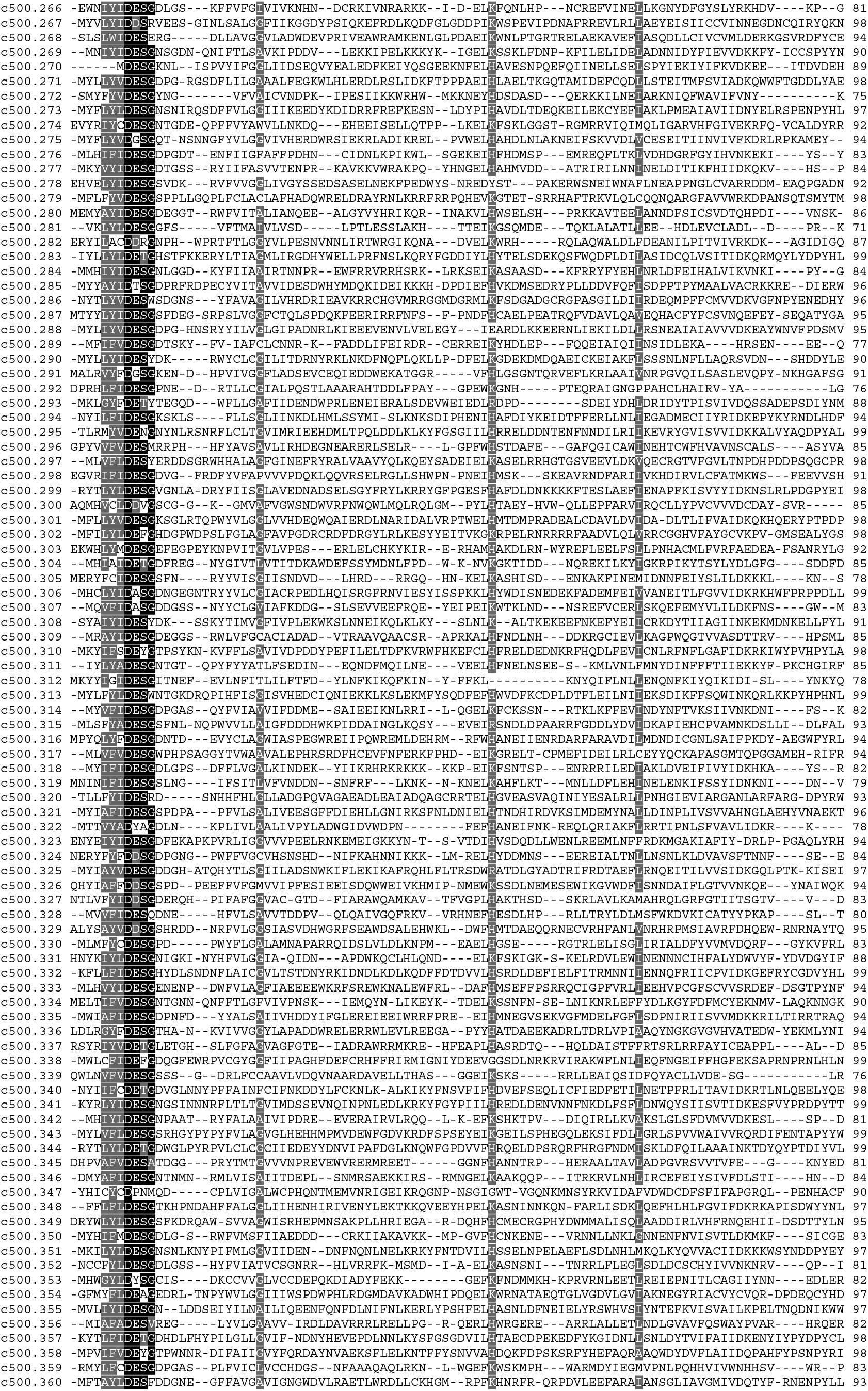
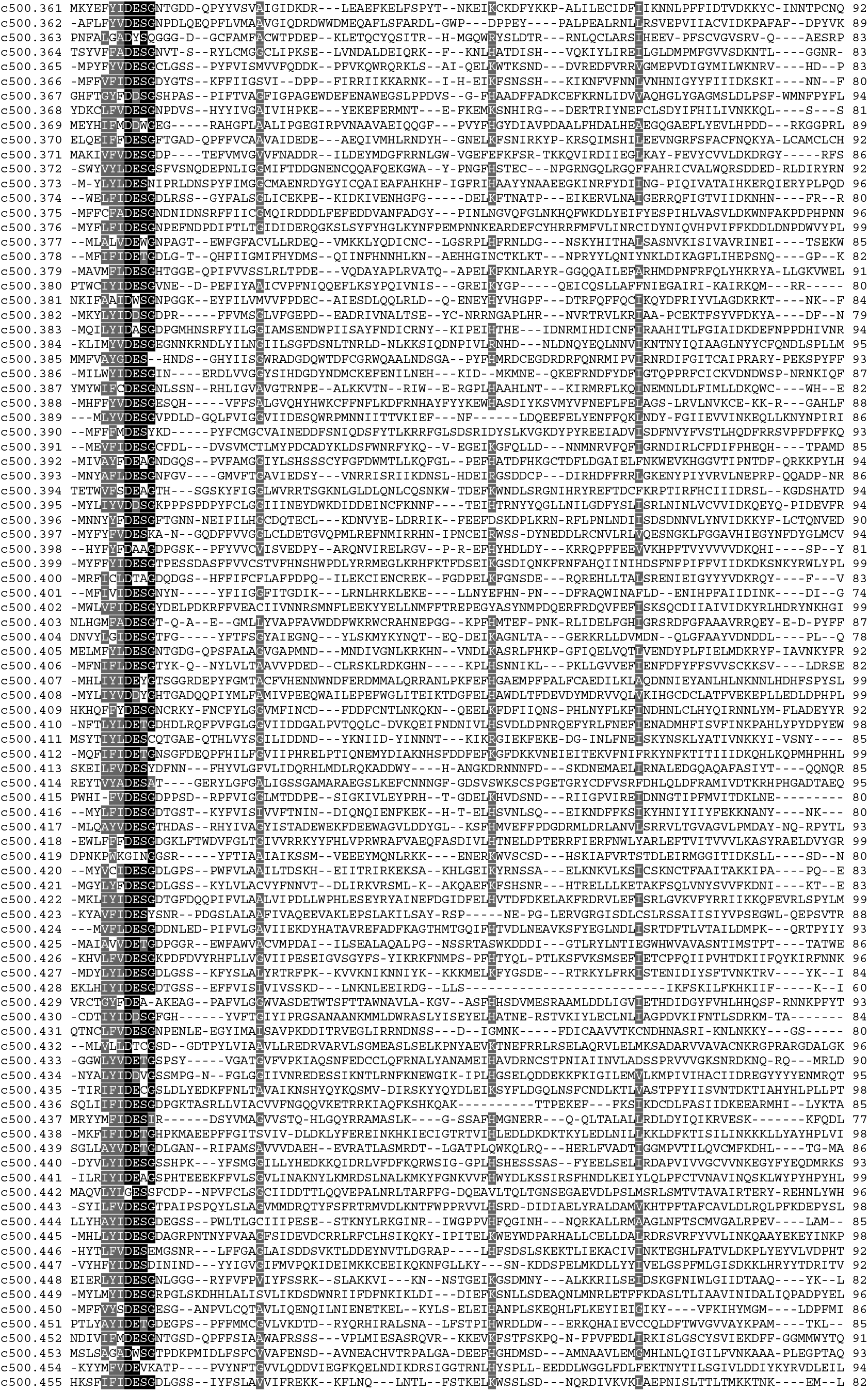
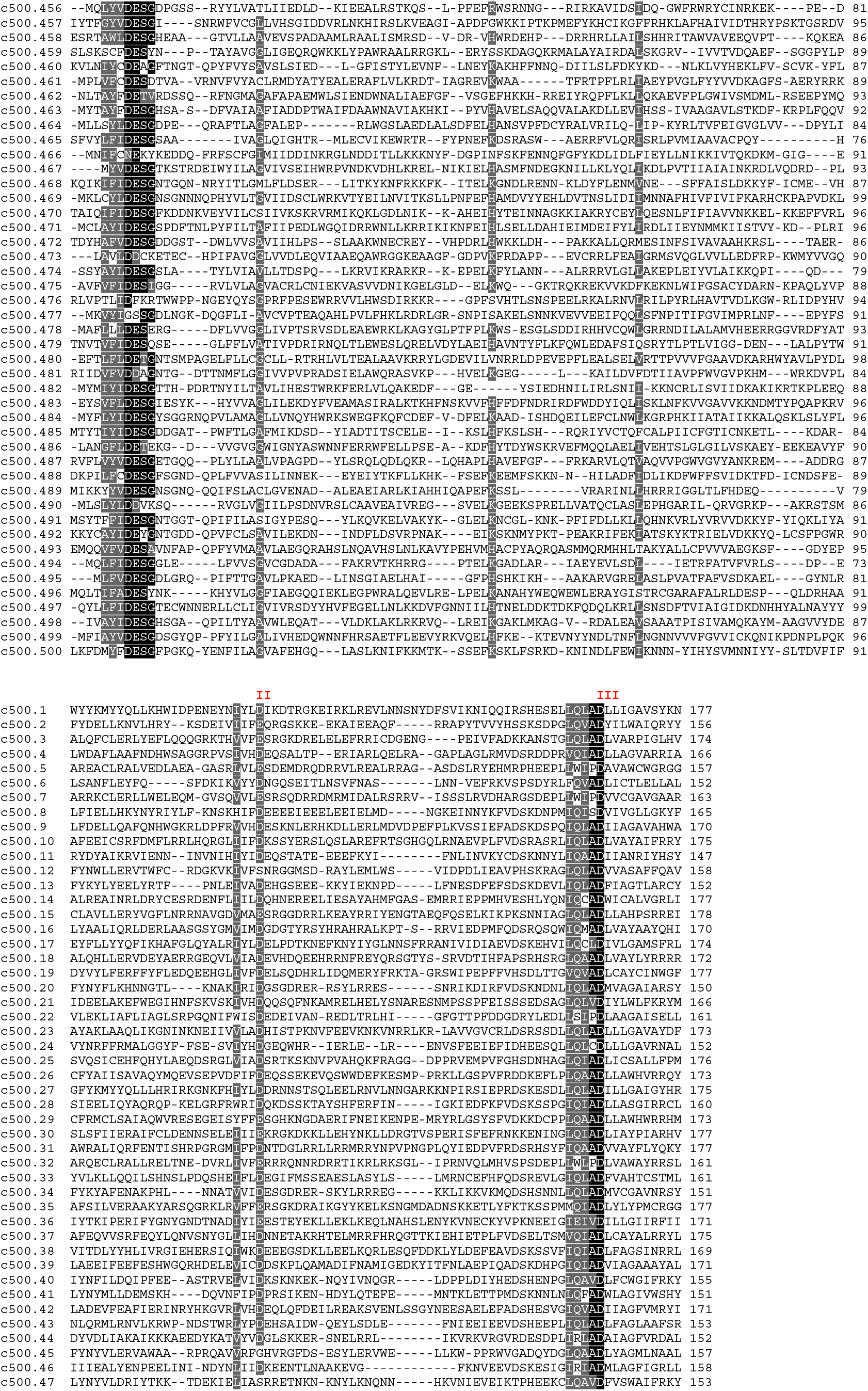
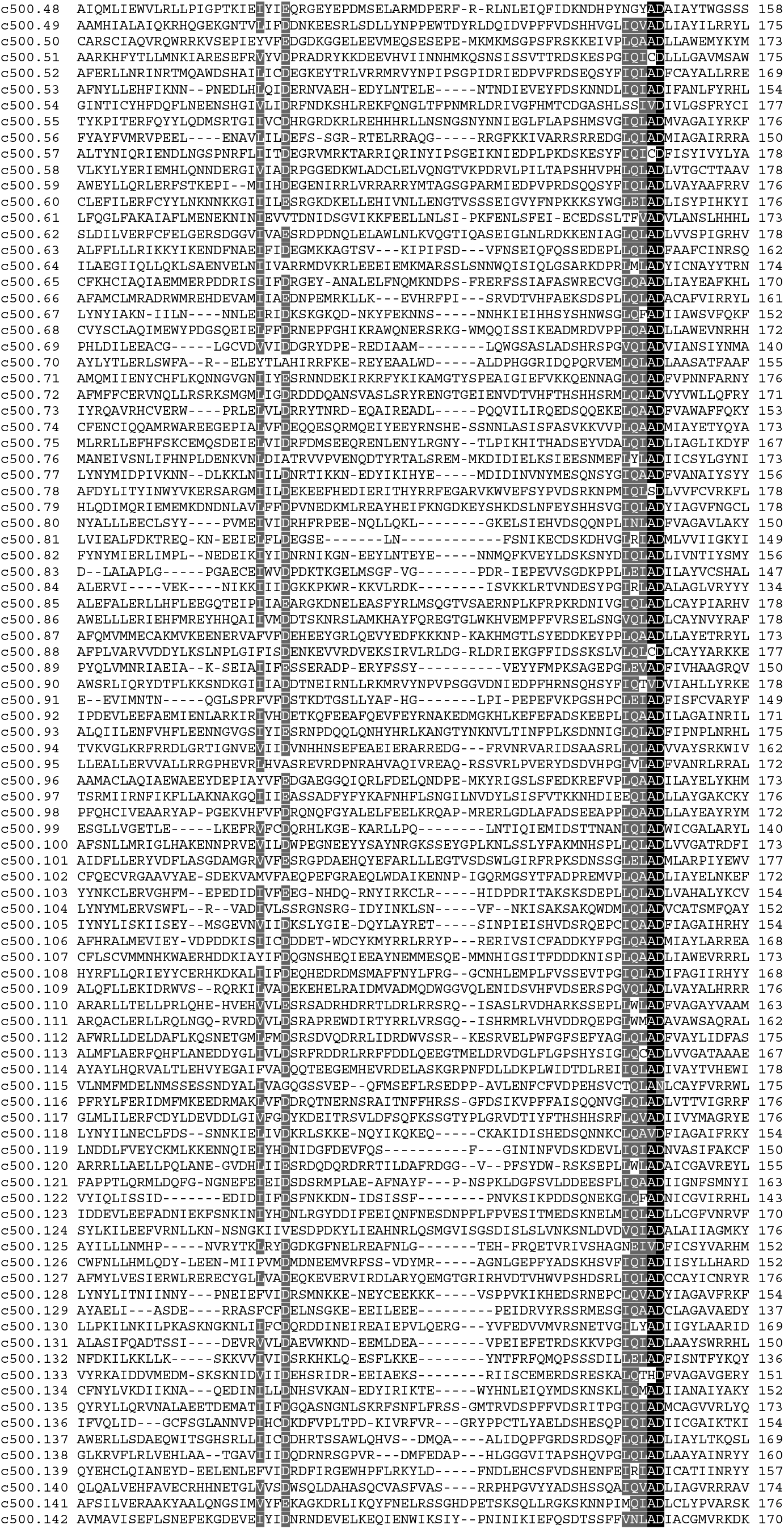
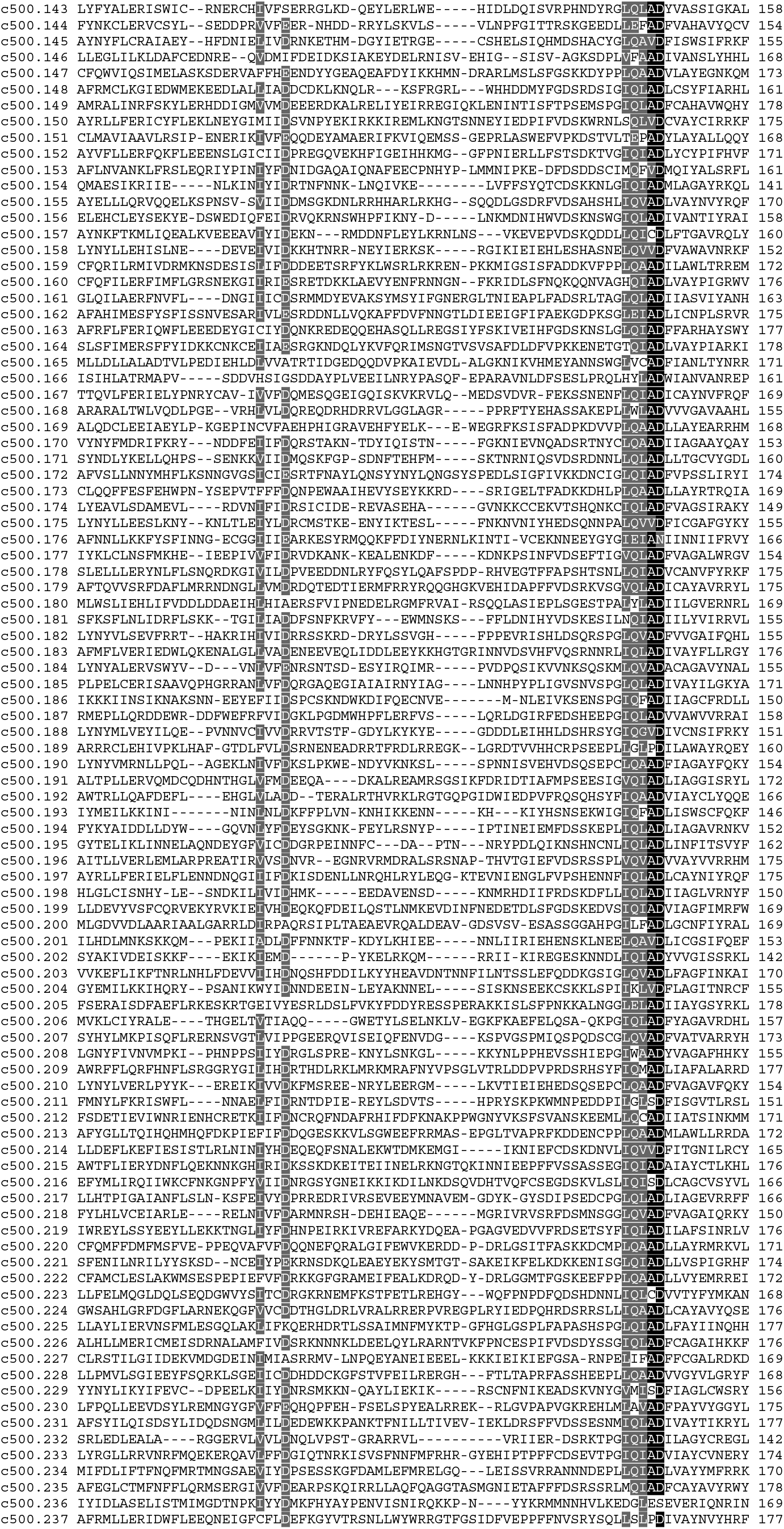
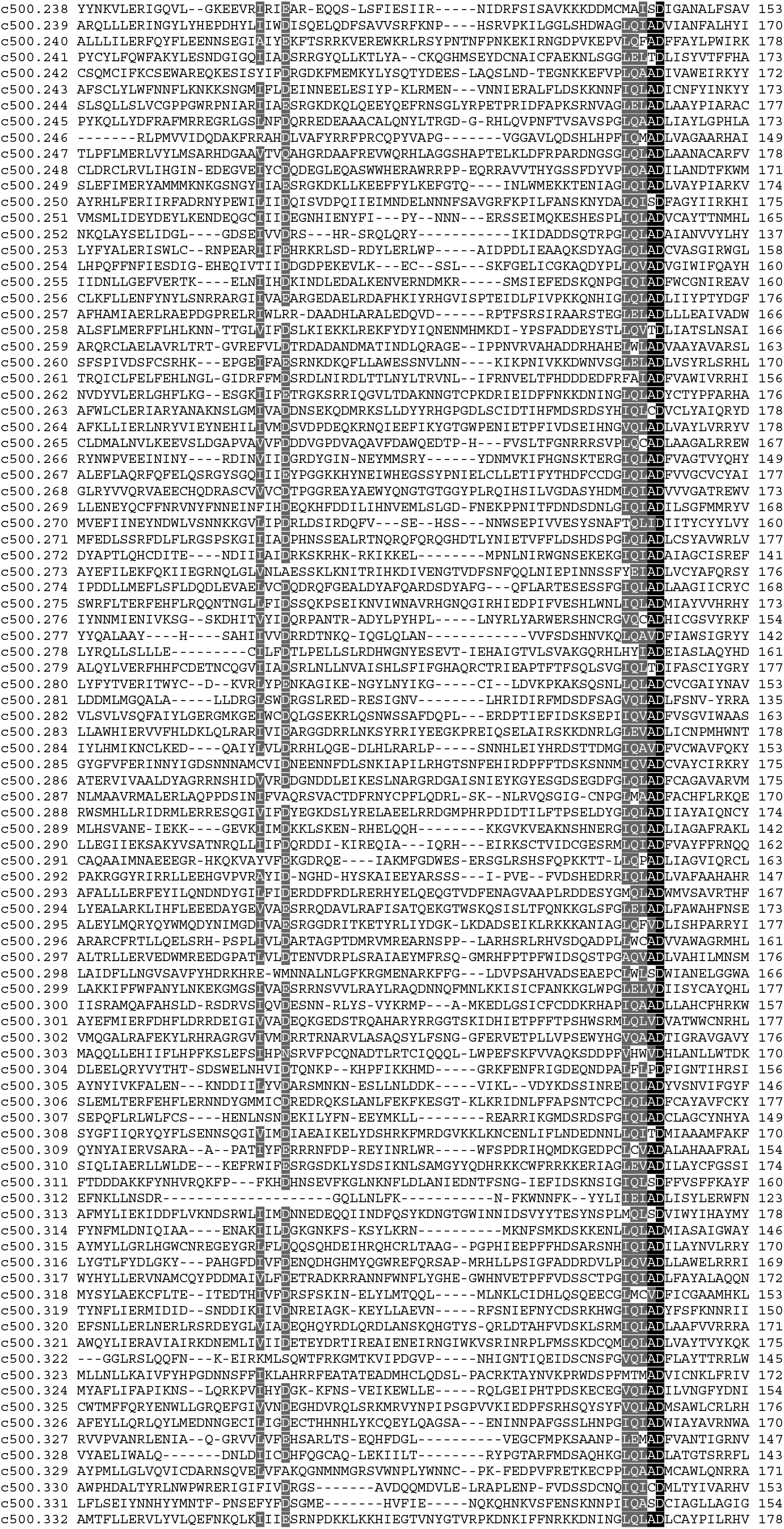
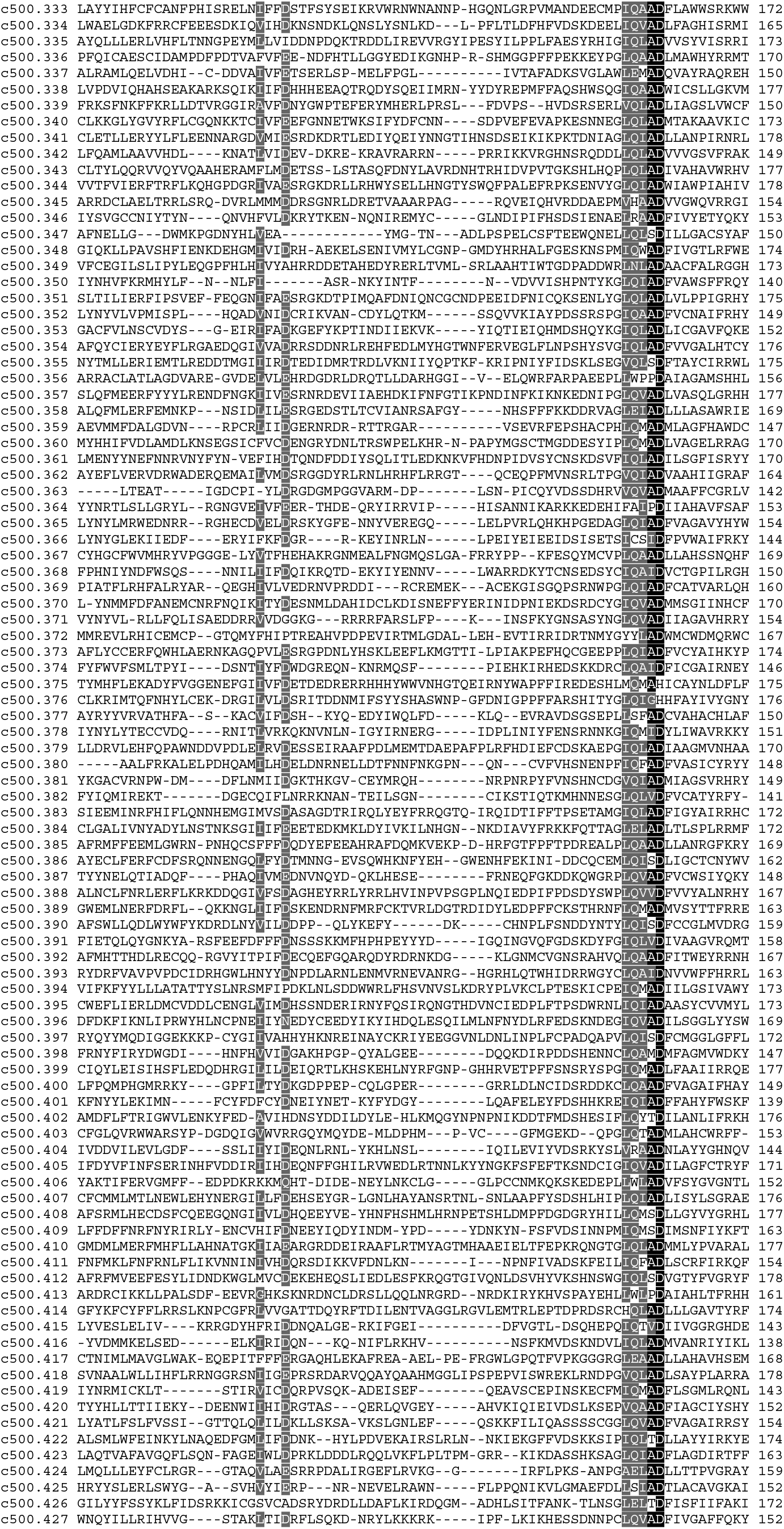
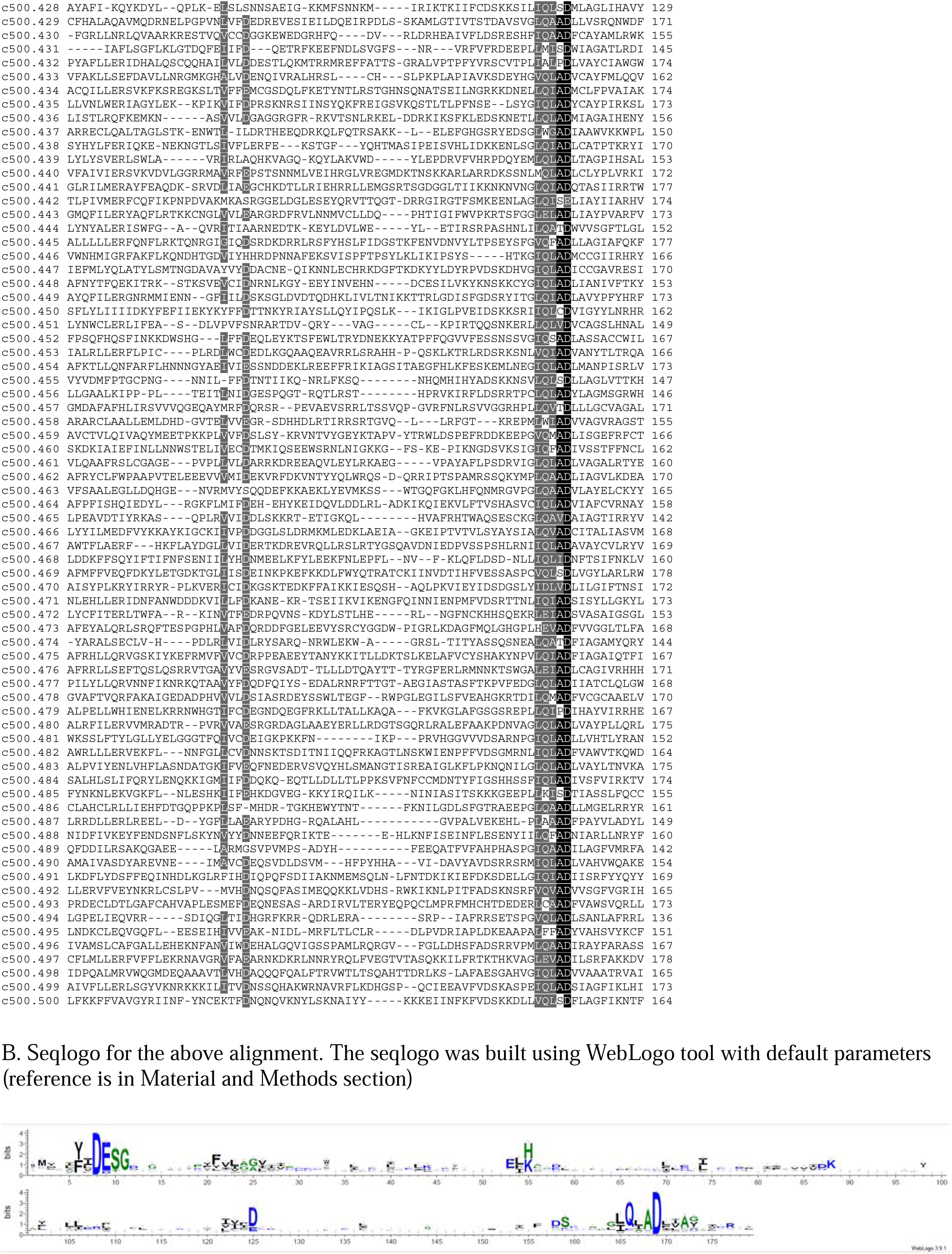
Sequence alignment and Seqlogo for DUF3800 protein family A. Alignment of 500 consensus sequences. These consensus sequences were built for each of 500 deep clades in the DUF3800 phylogenetic tree as described in Material and Method section. Alignment was colored with 80% similarity threshold with default amino acid groupings using https://www.bioinformatics.org/sms2/color_align_cons.html. Three RNase H catalytic sites are indicated by red (I, II and III). B. Seqlogo for the above alignment. The seqlogo was built using WebLogo tool with default parameters (reference is in Material and Methods section)

**Figure S2.**
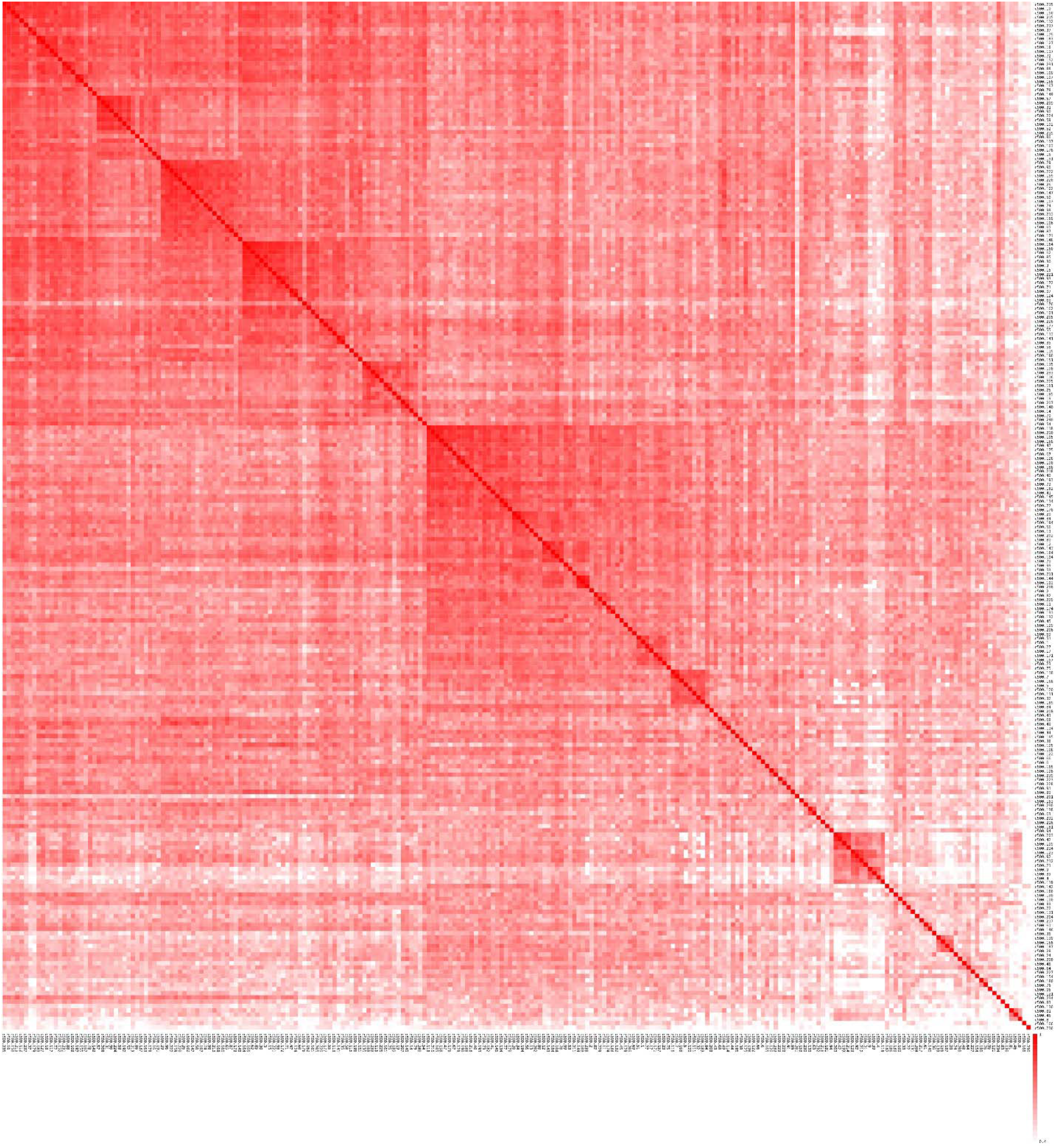
Structural comparison of DUF3800 domains The heat map shows the structural similarity (TM-score) between 240 DUF3800 structural models, representing clusters with 10 or more sequences. Alignments, structures and comparisons are available in Dataset 5.

**Figure S3.**
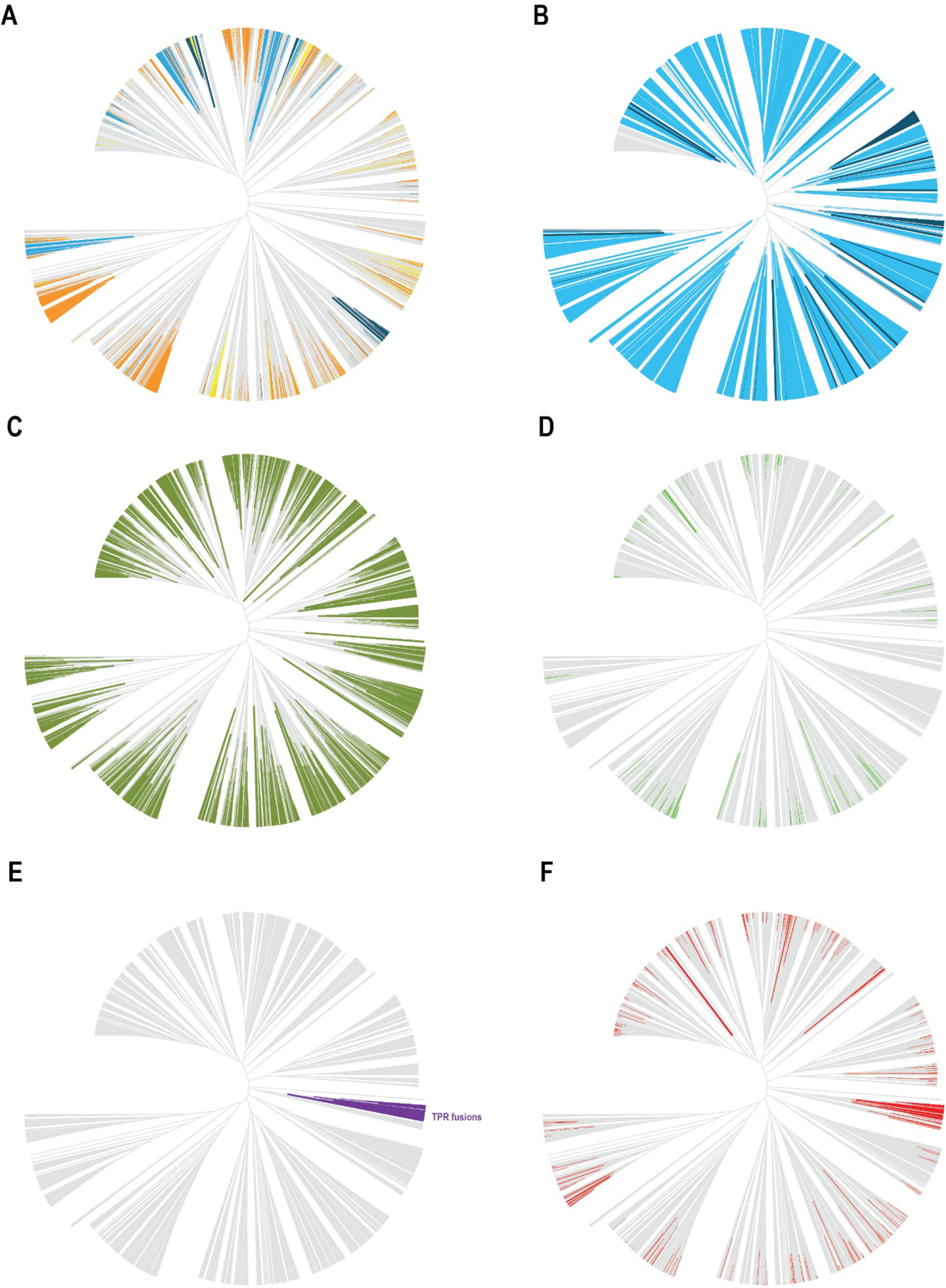
Selected features mapped onto DUF3800 tree. A. Map of DUF3800-I..XI RNA based on Rfam database scan and predicted in this work conserved regions (putative ncRNA): sky blue – DUF3800 downstream, dark blue DUF3800 upstream (if both up- and downstream ncRNA were identified only downstream is shown), orange – conserved intergenic regions downstream; yellow – conserved intergenic regions upstream (downstream region had a preference). Complete information is provided in Table S1. B. Space to code for RNA gene (90 nucleotides minimum): sky blue – downstream, dark blue -upstream. C. Presence of known defense genes (green) in the 10 genes vicinity of respective DUF3800 gene. Defense genes were identified using DefenseFinder. Details are in Table S1. D. DUF3800 imbedded in the RM type I operons (green). E. TPR domain fusions (purple). F. Presence of components of CRISPR-Cas systems in 10 genes vicinity of respective DUF3800 gene (red).

**Figure S4.**
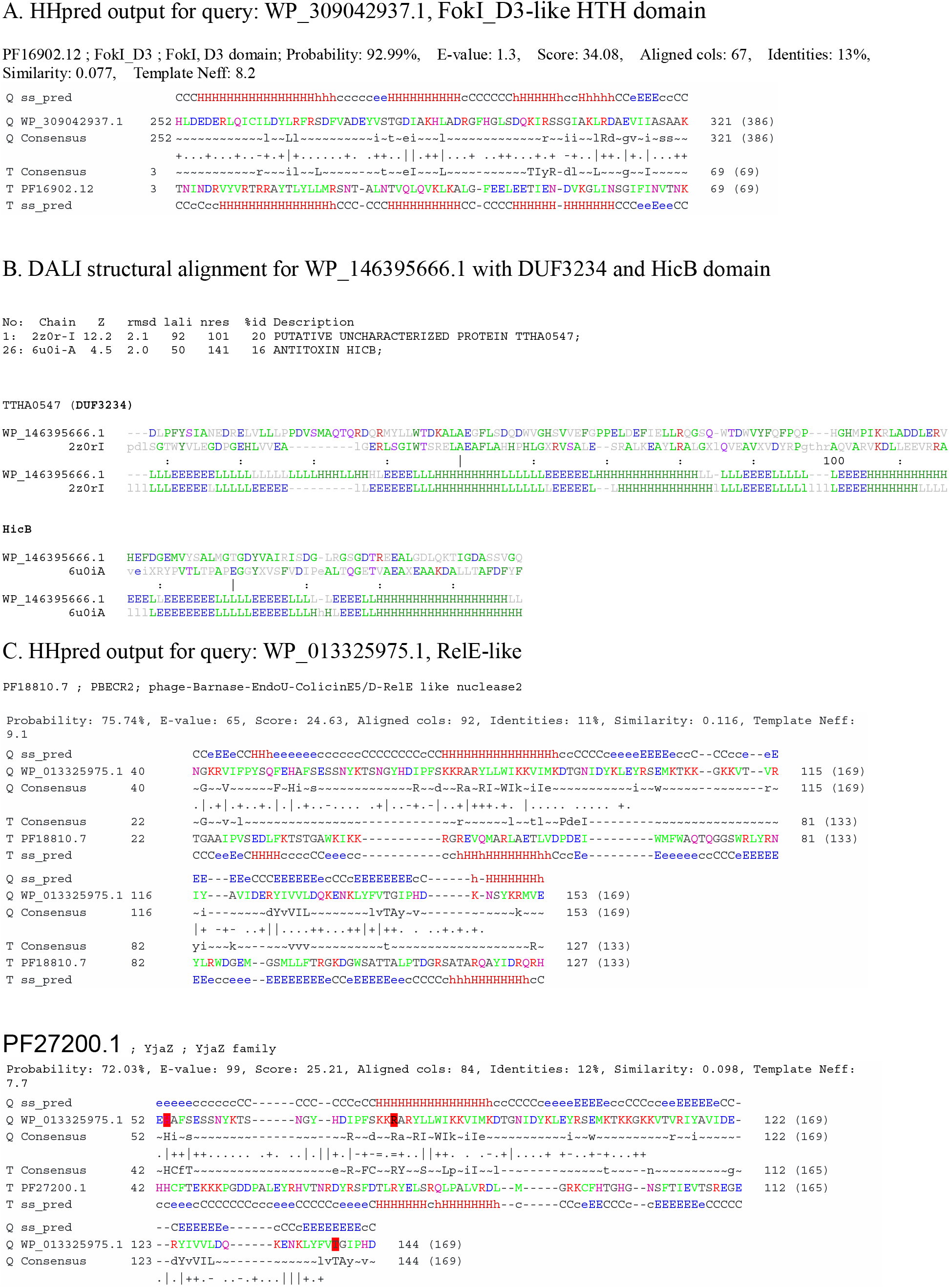
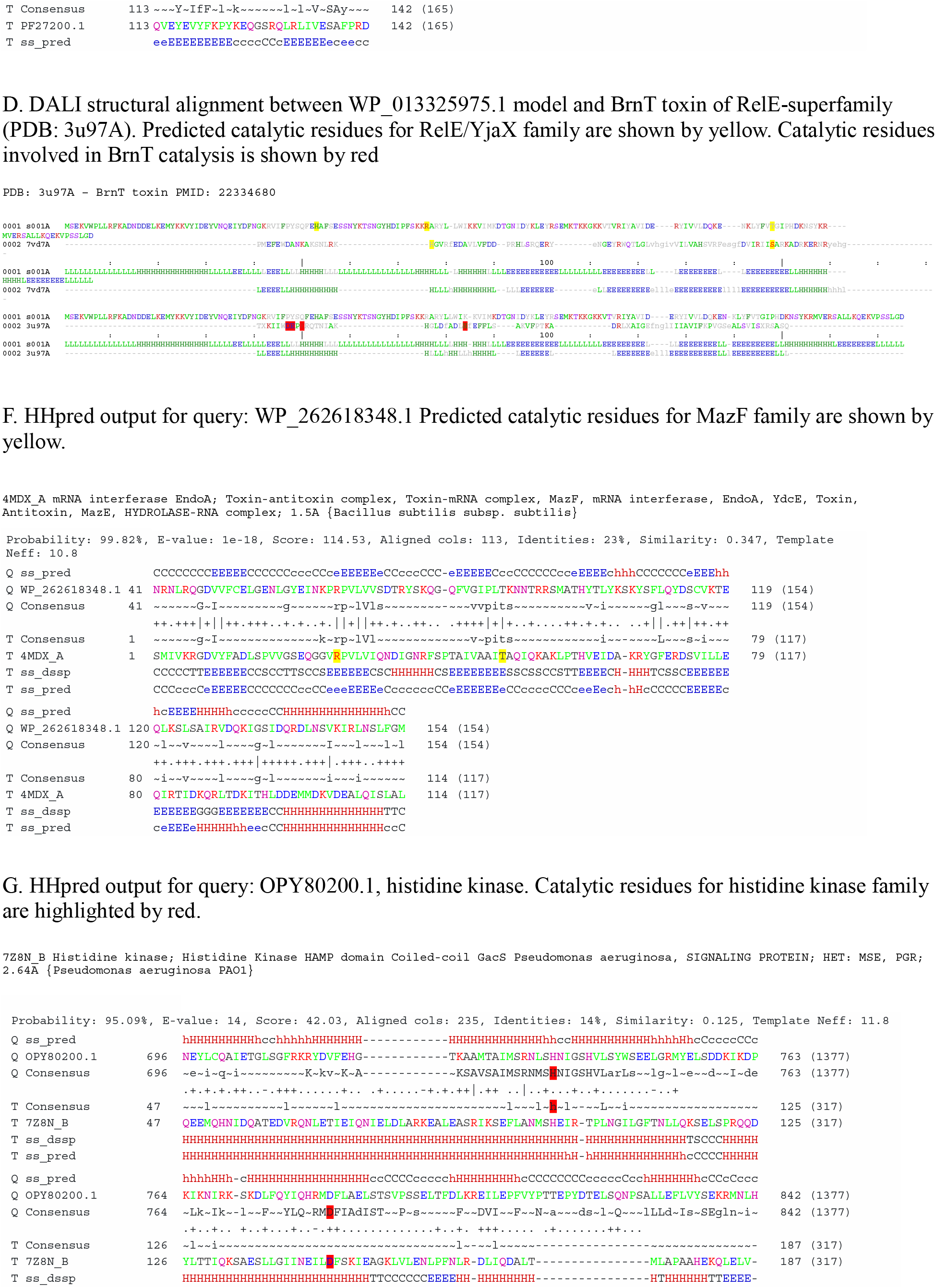
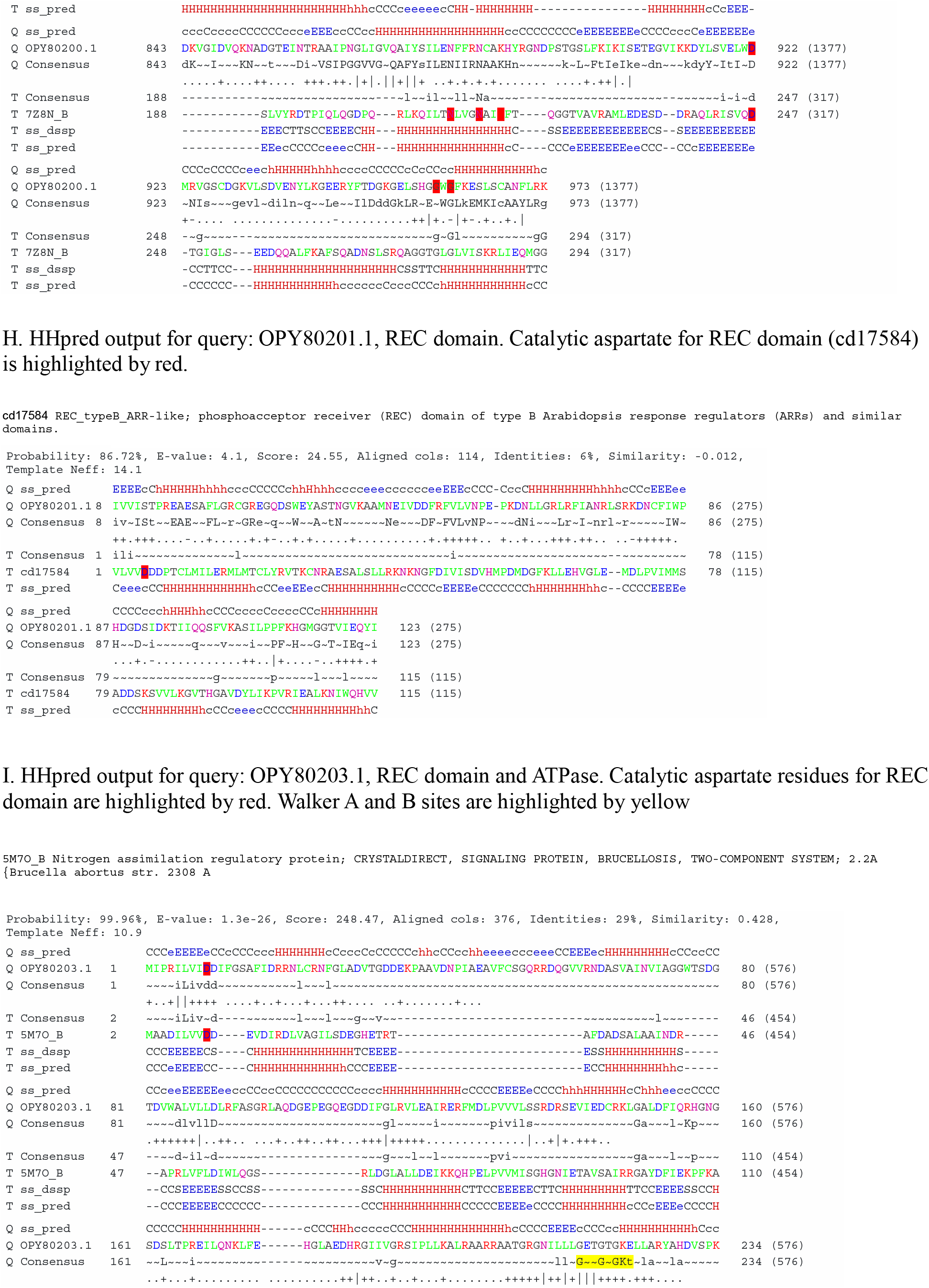
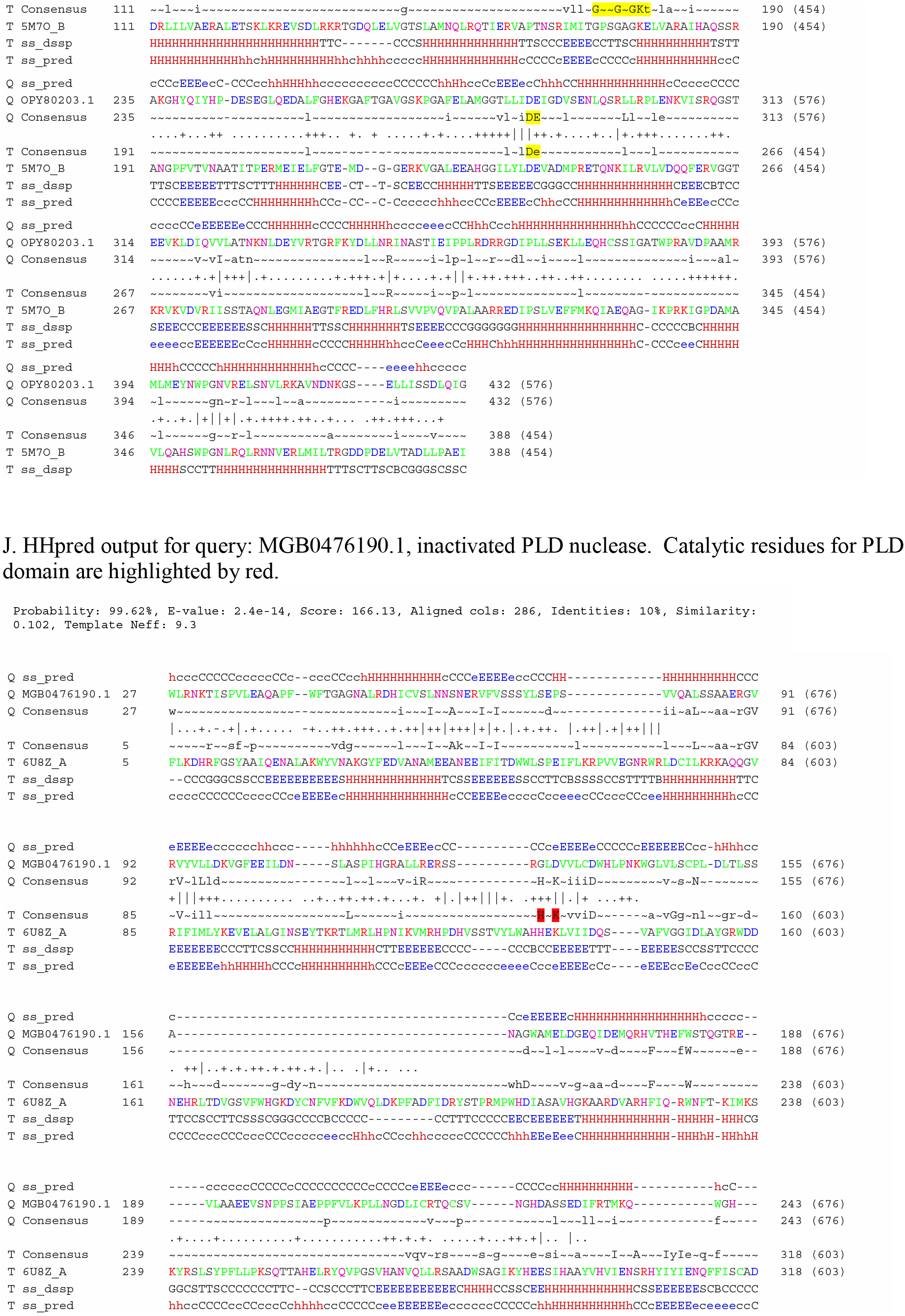
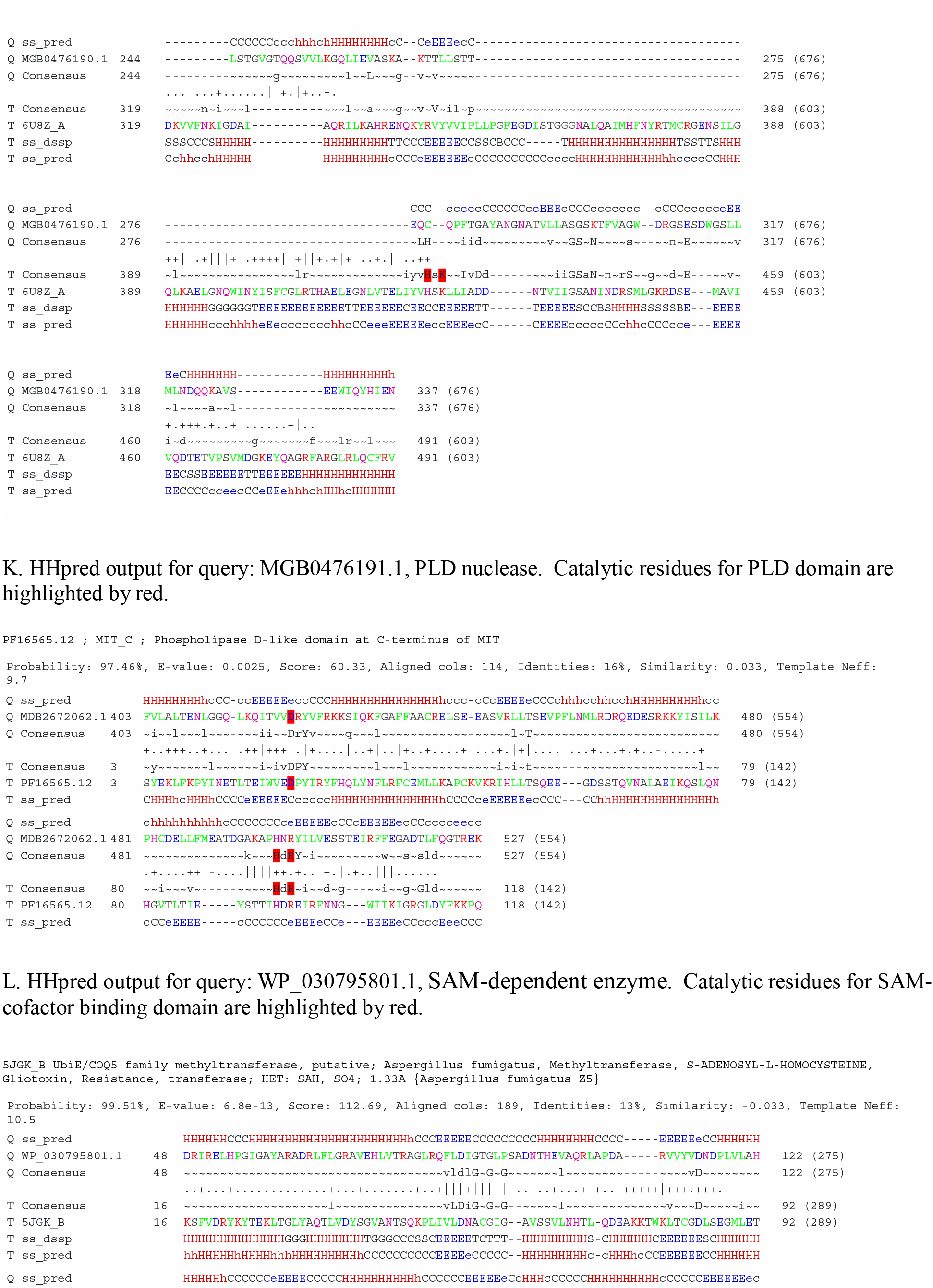
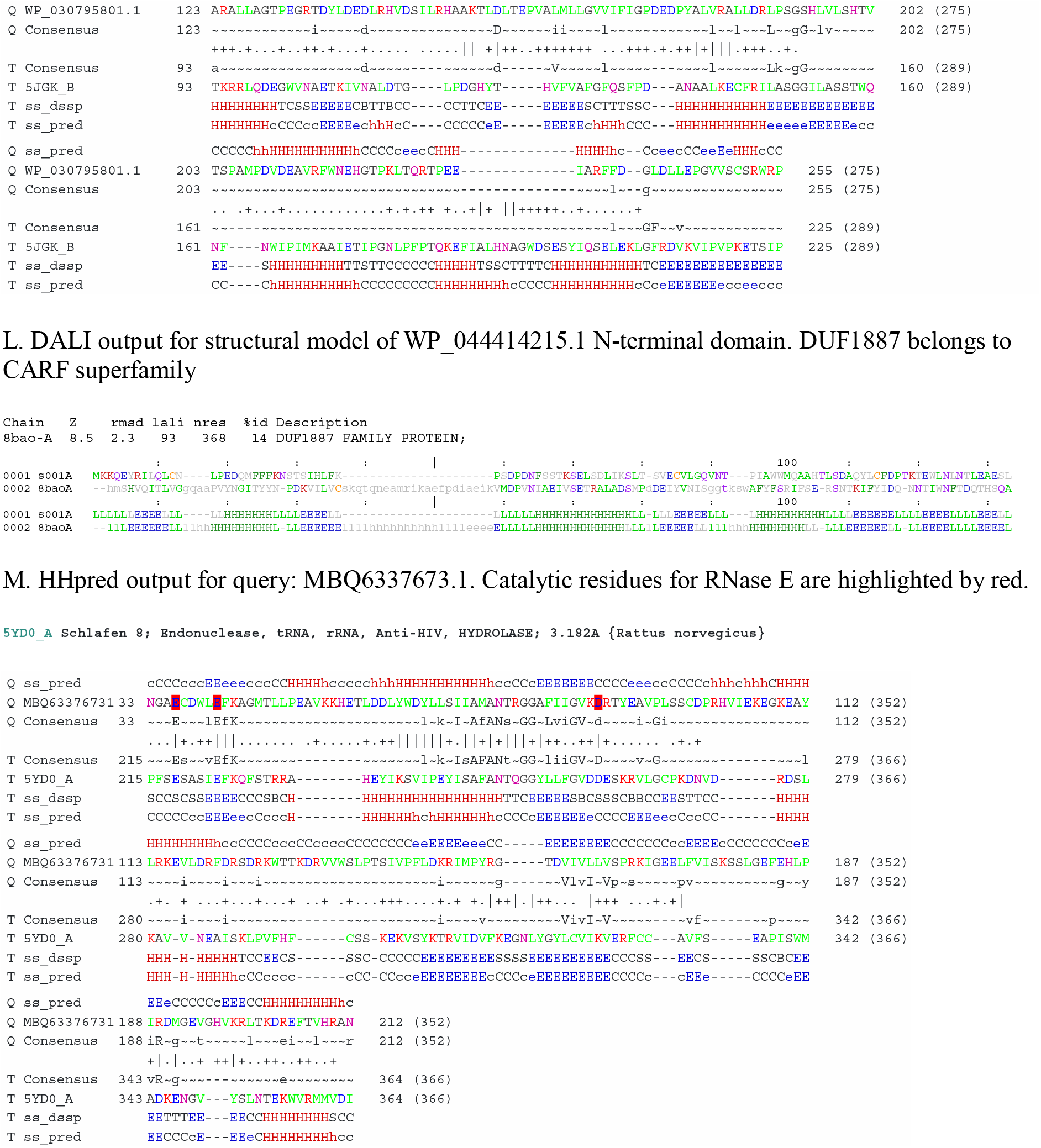
Sequence analysis of proteins described in the text. A. HHpred output for query: WP_309042937.1, FokI_D3-like HTH domain B. DALI structural alignment for WP_146395666.1 with DUF3234 and HicB domain. C. HHpred output for query: WP_013325975.1, BECR nuclease. D. DALI structural alignment between WP_013325975.1 model and BrnT toxin of BECR-superfamily (PDB: 3u97A). Predicted catalytic residues for BECR/YjaX family are shown by yellow. Catalytic residues involved in BrnT catalysis is shown by red. F. HHpred output for query: WP_262618348.1 Predicted catalytic residues for MazF family are shown by yellow. G. HHpred output for query: OPY80200.1, histidine kinase. Catalytic residues for histidine kinase family are highlighted by red. H. HHpred output for query: OPY80201.1, REC domain. Catalytic aspartate for REC domain (cd17584) is highlighted by red. I. HHpred output for query: OPY80203.1, REC domain and ATPase. Catalytic aspartate residues for REC domain are highlighted by red. Walker A and B sites are highlighted by yellow. J. HHpred output for query: MGB0476190.1, inactivated PLD nuclease. Catalytic residues for PLD domain are highlighted by red. K. HHpred output for query: MGB0476191.1, PLD nuclease. Catalytic residues for PLD domain are highlighted by red. L. HHpred output for query: WP_030795801.1, SAM-dependent enzyme. Catalytic residues for SAM-cofactor binding domain are highlighted by red. M. DALI output for structural model of WP_044414215.1 N-terminal domain. DUF1887 belongs to CARF superfamily. N. HHpred output for query: MBQ6337673.1, Schlafen/RNase E. Catalytic residues for RNase E are highlighted by red.

**Figure S5.**
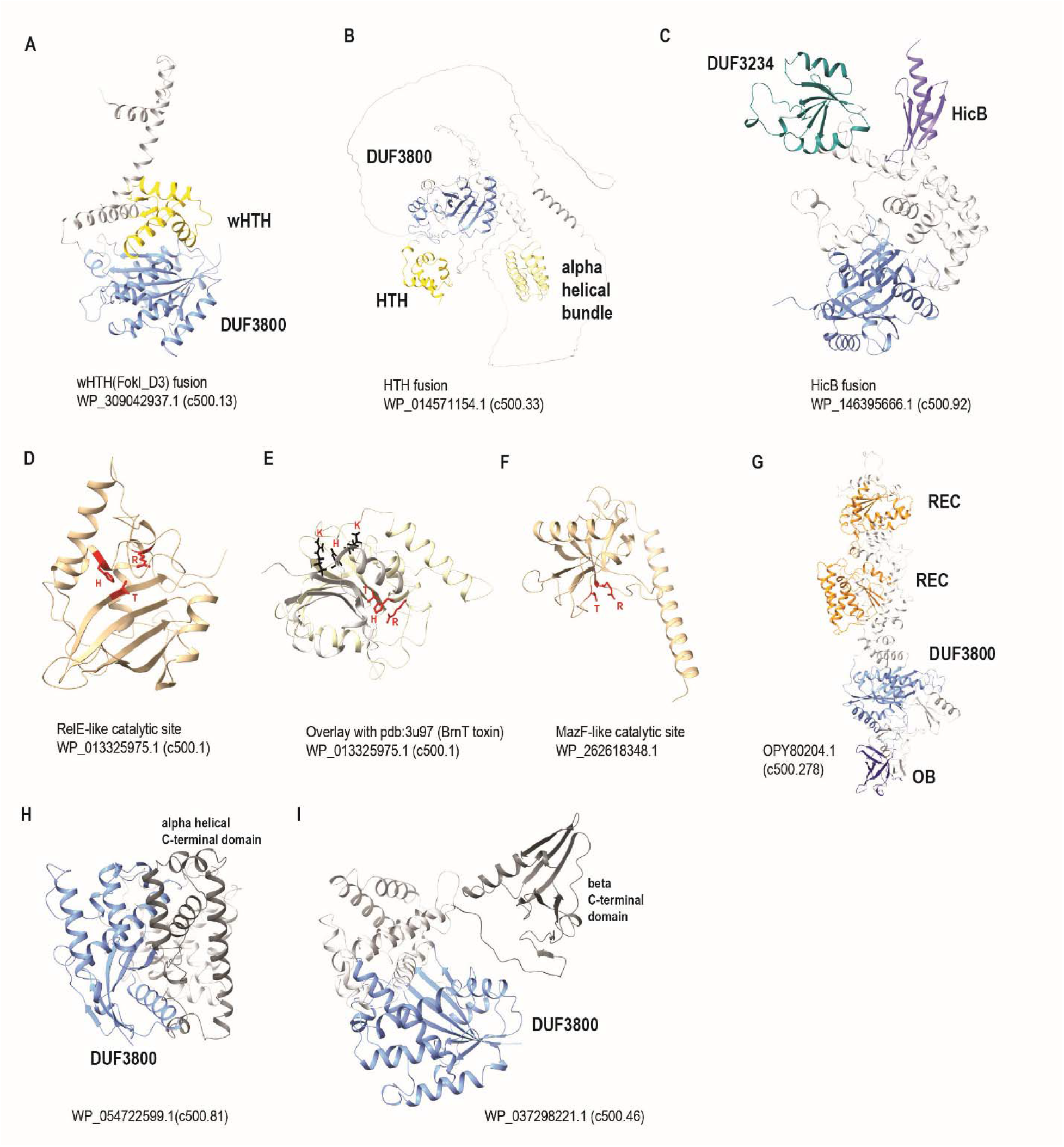
Structural models and comparison of the predicted structure of selected proteins A. Model of DUF3800 protein fused to FokI-D3 HTH domain. DUF3800 is shown by ski blue and HTH – by bright yellow. B. Model of DUF3800 protein fused to HTH domain and helical bundle domain. DUF3800 is shown by ski blue, helical bundle domain – pale yellow, HTH – by bright yellow. C. Model of a predicted RNase of BECR superfamily. Putative catalytic residues are shown by red. D. Overlay of BECR nuclease with pdb:3u97 (BrnT toxin). Putative catalytic residues of BECR-nuclease are shown by red; catalytic residues of BrnT toxin are shown by black. F. Model of MazF-like predicted RNase. Catalytic residues are shown by red. G. Model of DUF3800 protein fused to REC and OB domains. Rec domain are shown by orange and OB domain is shown by dark blue. H. HHpred output for query: OPY80201.1, REC domain. Catalytic aspartate for REC domain (cd17584) is highlighted by red. I. HHpred output for query: OPY80203.1, REC domain and ATPase. Catalytic aspartate residues for REC domain are highlighted by red. Walker A and B sites are highlighted by yellow. G. Model of DUF3800 protein fused to unknown alpha-helical domain (dark gray). H. Model of DUF3800 protein fused to unknown beta stranded domain (dark gray). Accession numbers for all models are indicated.

**Figure S6.**
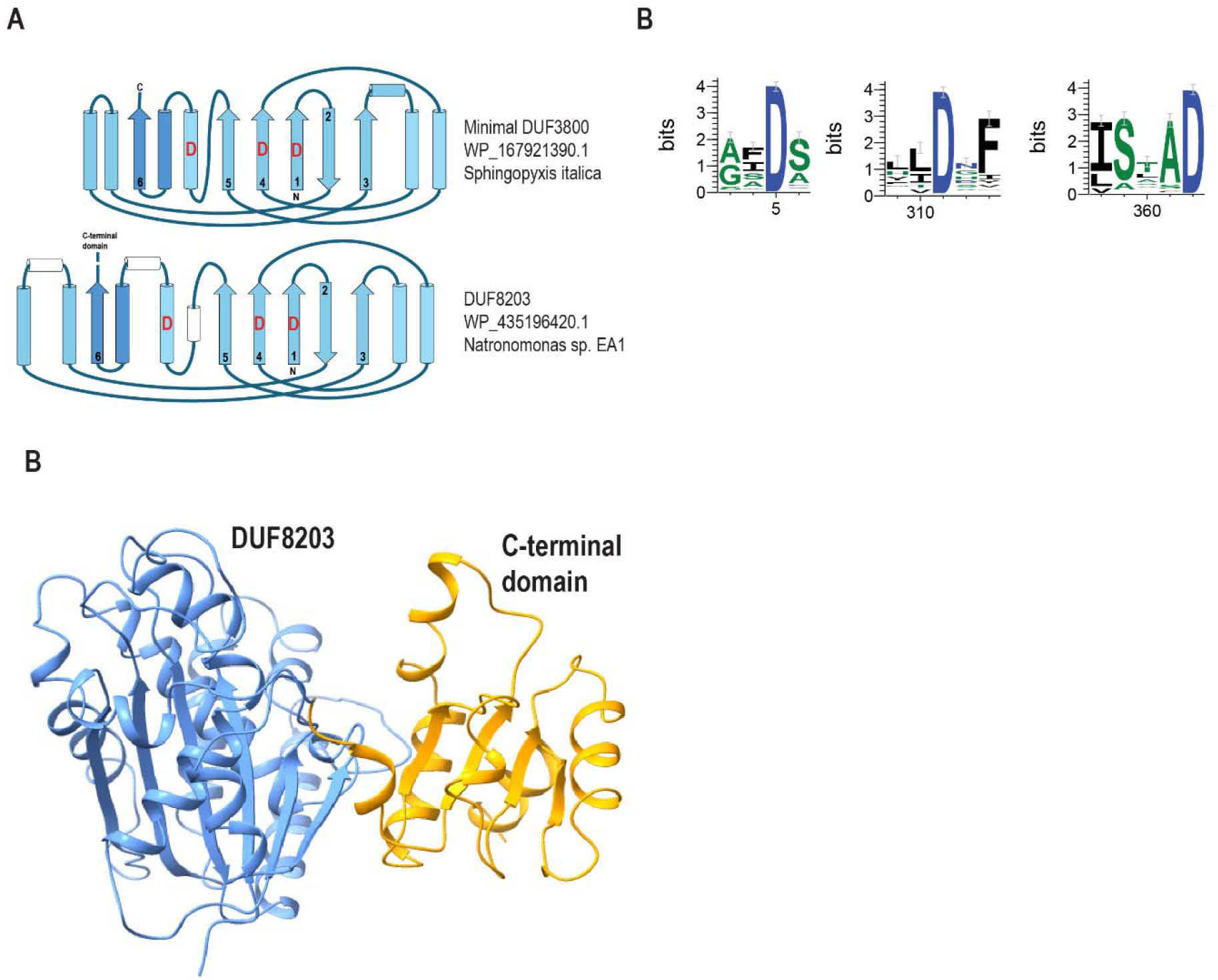
Analysis of DUF8203 family A. Topology diagram comparing DUF3800 and DUF3800 models. Beta strands are numbered. Catalytic aspartates are shown by red. B. Seqlogo based on multiple alignment of 152 DUF8203 sequences. Only conserved motifs that include catalytic aspartates are shown. Complete alignment is available in the Dataset 2. C. AF3 Model of DUF8203 protein. RNase H domain is shown by sky blue and C-terminal domain by yellow.

**Table S1.** DUF3800 protein family analysis

The table contains complete list of DUF3800 proteins included in the tree and metadata supporting Figure 1A,1G,2A,4A and Supplementary Figure S2.

**Table S2.** DUF8203 protein family analysis

The table contains complete list of DUF8203 proteins and metadata including taxonomy, up- and downstream region length, and presence of defense systems

**Table S3.** Identified and predicted DUF3800-associated ncRNAs

The data includes list of identified DUF3800-I-XI RNAs and conserved up- and downstream regions identified in this work. For predicted conserved region RNA folding statistics, length, number of sequences in each cluster and consensus sequence for each region are provided. Same data is shown for DUF8203. Also, the percent of potential ncRNA identified for each clade in the tree is shown.

**Table S4.** Direction analysis of DUF3800

The list of familied found to be codirected with DUF3800, either up- or downstream is shown. The families are sorted by the number of occurrences in directions.

**Table S5.** Proteins tested in this study

The list of accession numbers for the WARD systems tested. Annotation of genes in closest vicinity of the studied genes.

**Table S6.** The bacterial strains, plasmids and phages used in this study

The table contains a complete list of bacterial strains and plasmids tested in this study as well as a list of the bacteriophage and how they were maintained.

**Table S7.** WARD phage assay screen data

This table contains the numbers underlying the heatmap shown in Figure 5B.

**Table S8.** Membrane permeability over the course of infection

This table contains the complete dataset of underlying Figure 6C and 6F monitoring the levels of propidium iodide over the course of infection.

## Datasets

**Dataset 1. Multiple alignments for 500 clades in DUF3800 tree.**

**Dataset 2. Multiple alignment for DUF8203 protein family.**

**Dataset 3. DUF3800 proteins identified in complete genome database.**

**Dataset 4. DUF3800 tree in Newick format and corresponding alignment.**

**Dataset 5. Structure prediction and comparison of DUF3800 domains.**

**Dataset 6. Alignments of conserved intergenic regions identified in this work.**

**Dataset 7. Genomic neighborhoods of DUF3800 genes.**

